# Oceanic cyanobacterial photosynthesis is negatively affected by viral NblA proteins

**DOI:** 10.1101/2024.11.10.622831

**Authors:** Omer Nadel, Rawad Hanna, Andrey Rozenberg, Dror Shitrit, Ran Tahan, Irena Pekarsky, Oded Béjà, Oded Kleifeld, Debbie Lindell

## Abstract

Marine picocyanobacteria are abundant photosynthetic organisms of global importance. They coexist in the ocean with cyanophages, viruses that infect cyanobacteria. Cyanophages carry many auxiliary metabolic genes acquired from their hosts that are thought to redirect host metabolism for the phage’s benefit^1–5^. One such gene is *nblA* which is present in multiple cyanophage families^2,6–8^. Under nutrient deprivation the cyanobacterial NblA is responsible for inducing proteolytic degradation of the phycobilisome^9–11^, the large cyanobacterial photosynthetic light harvesting complex. This increases the pool of amino acids available for essential tasks^11^, serving as a survival mechanism^12^. Ectopic expression of different cyanophage *nblA* genes results in host pigment protein degradation^9,6^. However, the benefit of the cyanophage-encoded NblA for the cyanophage and the broader impact on the host are unknown. Here, using a recently developed genetic manipulation system for marine cyanophages^14^, we reveal that cyanophage NblA significantly accelerates the cyanophage infection cycle, directs degradation of the host phycobilisome and other proteins and reduces host photosynthetic light harvesting efficiency. Metagenomic analysis revealed that cyanophages carrying *nblA* are widespread in the oceans and compose 35% and 65% of oceanic T7-like cyanophages in the surface and deep photic zones, respectively. Our results show a large benefit of the *nblA* gene to the cyanophage while exerting a negative effect on the host photosynthetic apparatus and host photosynthesis. These findings suggest that cyanophage *nblA* has an adverse global impact on light harvesting by oceanic picocyanobacteria.

---

The light-harvesting antennae of most cyanobacteria are composed of phycobilisomes, gigantic membrane-associated soluble supercomplexes, located on the exterior side of the thylakoid membrane^9,15^. Cyanobacteria capture and funnel solar energy through their phycobilisomes towards the photosynthetic reaction centers^16^ where the primary energy conversion reactions of photosynthesis occur. Typical phycobilisomes of marine *Synechococcus* are built of multiple subunits that absorb light at different wavelengths, including an allophycocyanin (APC) core and peripheral rods composed of disks of different phycobiliproteins, phycocyanin (PC) and phycoerythrin-I and -II (PEI, PEII) as well as linker proteins that connect rod subunits to each other and to the core and attach the phycobilisome to the thylakoid membrane^17,18^. Each phycobiliprotein is composed of α and β subunits that form heterodimers^10^. These proteins bind the pigment chromophores phycocyanobilin, phycoerythrobilin (PEB) and phycourobilin (PUB), which absorb light at different wavelengths^19^. Due to their large size, phycobilisomes can comprise up to half of the soluble protein content of a cell^15^. *Prochlorococcus* is the most abundant photosynthetic organism on Earth^20^. The vast majority of *Prochlorococcus* lineages have a chlorophyll-based transmembranal light harvesting complex rather than the typical phycobilisome^21–23^. *Prochlorococcus* does, however, code for one or two subunits of phycoerythrin (PE)^24,25^, found at low levels in cells^26^. Interestingly, some low-light adapted *Prochlorococcus* lineages from anoxic marine zones possess both chlorophyll-based and phycobilisome light harvesting complexes^20^.

## Cyanophage NblA is beneficial for infection

Evidence collected over the past two decades suggests that cyanophages possess host-like auxiliary metabolic genes that are thought to influence key processes in host metabolism during infection and enhance phage replication^1–4^. These include genes that are thought to benefit the phage by maintaining essential cellular metabolic processes as well as those that impair host activities to redirect resources towards phage progeny production^3,4^. The *nblA* gene codes for a proteolysis adaptor protein found in multiple cyanophage families^2,6–8,13,27^, and is particularly prevalent in T7-like cyanophages (family *Autographiviridae*). In nutrient deprived cyanobacteria, the cyanobacterial NblA is responsible for inducing degradation of the phycobilisome complex, resulting in clear loss of pigmentation^9–11^. Here, we use S-TIP37, a typical T7-like cyanophage isolated from the Red Sea that infects the open ocean *Synechococcus* sp. strain WH8109^28^, as our model system to study cyanophage-encoded *nblA*. We began by examining whether the cyanophage *nblA* gene is transcribed and translated during the infection process. We found high *nblA* transcript and protein levels from two hours post-infection onwards (Extended Data Fig. 1). This expression pattern was distinct to that of a host copy of the *nblA* gene which displayed fairly consistent transcript and protein levels (Extended Data Fig. 1). The phage *nblA* expression pattern was similar to that of the phage DNA polymerase gene, and is thus expressed with phage genes involved in DNA metabolism and replication and prior to virion morphogenesis genes^29,30^.

Next, we assessed the influence of the *nblA* gene on cyanophage infection. For this we constructed *nblA* cyanophage deletion mutants (Δ*nblA*) using the REEP approach^14^ (see Methods). Infection of the host with the Δ*nblA* mutant phage revealed visibly less pigmentation loss than when the host cells were infected with the wild-type (WT) phage (Fig. 1a), indicating a clear phenotype for the Δ*nblA* mutant during infection.

**Fig. 1.**
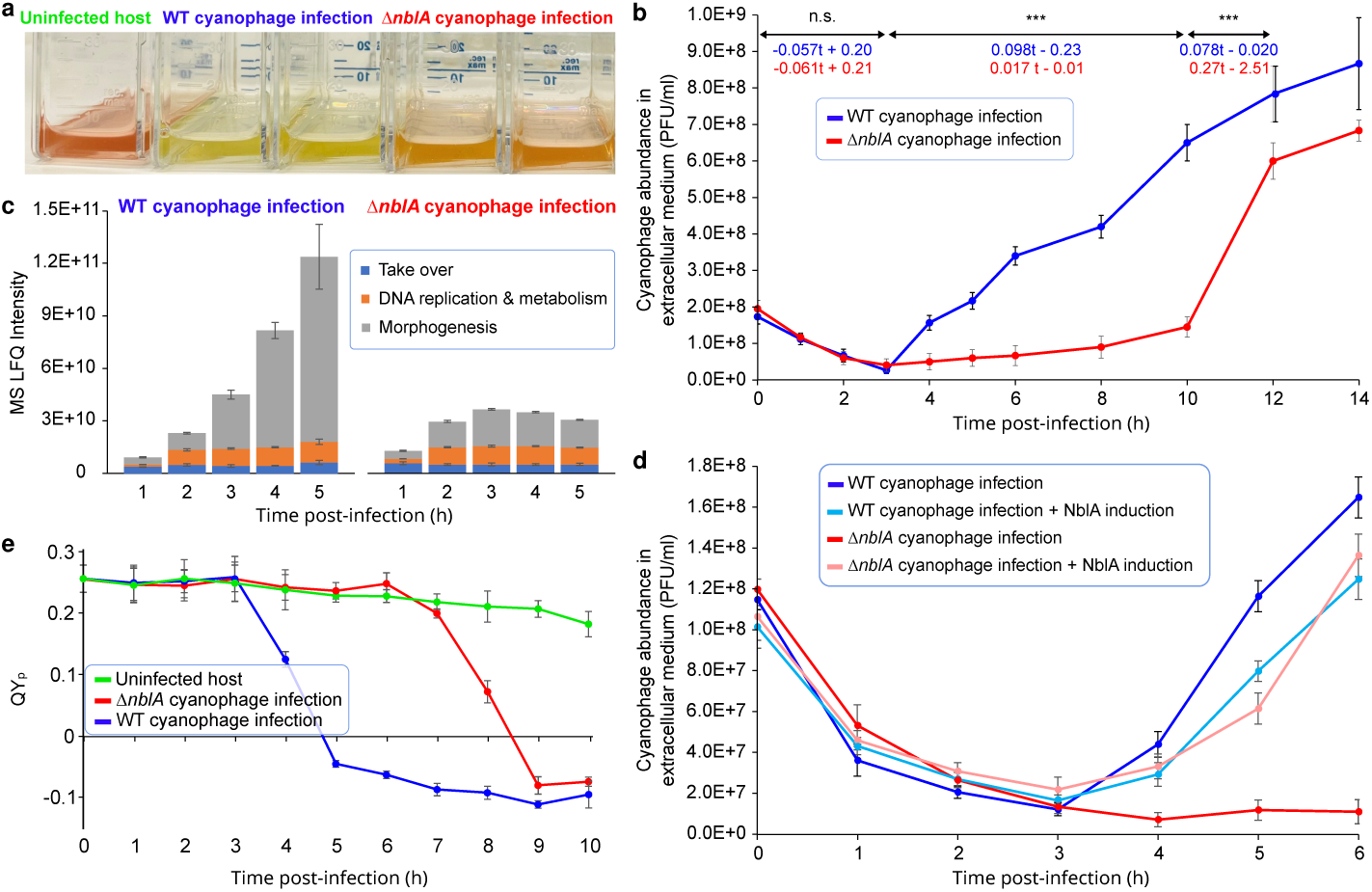
Influence of the *nblA* gene on S-TIP37 cyanophage infection dynamics. **a**, Culture pigmentation phenotypes in uninfected *Synechococcus* sp. strain WH8109, and in WH8109 6 h post-infection when infected with WT S-TIP37 or the Δ*nblA* mutant S-TIP37 cyanophage. **b**, Cyanophage growth curves of the WT and Δ*nblA* mutant cyanophages, *n*=3. **c**, Proteomic characterization of different functional groups of phage proteins with time after infection of the cyanobacterium with the WT (left) and Δ*nblA* (right) cyanophages. Linear models were fit for every functional group and a significant contribution of the factor “infection type” was found for the morphogenesis genes (FDR-adjusted *p*-value < 0.001). For further statistical details refer to Suppl. Data File 2, *n*=3. **d**, Cyanophage infection dynamics of the WT and Δ*nblA* mutant phages when infecting the wild-type host or the host ectopically expressing the cyanophage NblA, *n*=3. **e**, Apparent photochemical quantum yield of PSII (QY_p_) in cells infected with the WT or the *ΔnblA* cyanophages and in uninfected control cells after preferential excitation of the phycobilisome at 495 nm (the maximal absorbance of PEII). Infection was performed at a multiplicity of infection (MOI) of 5, *n*=3. For **b**, summary of the linear regression fits are provided for the three near-linear segments: formulas of the relative cyanophage abundance for WT (in blue) and *ΔnblA* cyanophages (in red) as a function of time are provided alongside the significance levels for the factor “infection type” (type II Wald *F* test): n.s. – non-significant, *** – FDR-adjusted p-value < 0.001, see Suppl. Data File 2 for further details. For **c**, genes are grouped into three clusters and the cluster names reflect functions of the majority of the corresponding genes: take over (genes STIP37_1-12 in GenBank record MH540083.1), DNA replication and metabolism (genes STIP37_13-23 and STIP37_55), and morphogenesis (genes STIP37_24-54). The *nblA* gene is located between genes STIP37_18 and STIP37_19. The plotted values in panels **b**-**e** correspond to averages and the whiskers reflect standard deviations.

To investigate the importance of the cyanophage NblA to the infection process, we performed phage growth curve experiments. The Δ*nblA* mutant demonstrated a substantially longer latent period (the period of infection prior to the first release of phage progeny from cells) of 8-10 h compared to the 3 h latent period of the WT^31^ (Fig. 1b). In addition, synthesis of phage proteins increased with time during infection with the WT phage (Fig. 1c, Suppl. Data File 1), whereas most phage protein production ceased after 3 h during infection with the Δ*nblA* mutant with significantly fewer virion morphogenesis proteins produced in the latter (Fig. 1c). These findings suggest a direct link between phage protein production and NblA expression and indicate that NblA is advantageous to the WT phage. Since cyanobacterial NblAs are known to degrade phycobilisomes during nutrient stress^9–11^, and to release amino acids critical for cell survival^11,12^, we hypothesize that the phage NblA has a similar role during infection but that the release of amino acids is used for building new virions.

The gene density in the region of *nblA* in the phage genome is high, thus to verify that the observed phenotype is a result of the lack of the *nblA* gene and not due to potential side-effects caused by the deletion (see Methods), we performed a rescue experiment. This was achieved by infecting *Synechococcus* sp. strain WH8109 cells that were ectopically expressing the cyanophage *nblA* gene under an inducible expression system^32,33^ with the Δ*nblA* mutant phage (Fig. 1d). Indeed, expression of the cyanophage NblA restored the timing of infection to that found for the WT cyanophage (Fig. 1d). The shorter latent period for the WT phage suggests that the *nblA* gene would improve phage fitness by allowing it to carry out more rounds of infection. Our findings confirm that the *nblA* gene plays an important function in cyanophage infection and that it likely enhances phage fitness.

Cyanophage NblA would be expected to impact host photosynthesis if it directs phycobilisome degradation since phycobilisomes transfer energy to the photochemical reaction centers of the photosystems (PSI or PSII)^16,34^. We assessed the photosynthetic performance of PSII by measuring fluorescence at room temperature after exciting cultures with blue light (495 nm) which preferentially excites the most peripheral disks of the phycobilisome rods consisting of PEII. The apparent photochemical quantum yield of PSII (QY_p_) was then calculated. While uninfected cells maintained a steady PSII quantum yield, infection with both the WT and Δ*nblA* mutant phages resulted in a considerable decline (Fig. 1e). Notably, this decline reached low levels by 8 h after infection with the Δ*nblA* mutant, a delay of 4 h relative to infection with the WT cyanophage. This indicates that the cyanophage NblA negatively impacts the host’s PSII photochemical quantum yield by 50% over the period of infection. This is likely to have a dramatic effect on the ability of the host cell to produce the energy and reducing equivalents necessary for many cellular processes. This diminished photosynthetic performance could be attributed to targeted phycobilisome degradation, accelerated cell lysis, or a combination of both factors.

## Cyanophage NblA causes degradation of host phycobilisomes

Cyanobacterial NblAs are known to be involved in phycobilisome degradation. To assess whether the cyanophage NblA has a similar effect on host phycobilisome integrity, we monitored whole-cell fluorescence emission spectra as a measure of the energy transfer pathway from phycobilisomes to PSII. Samples collected at different time points were excited at the wavelength for PUB, which binds mainly to the peripheral PEII disk of the phycobilisome, and the corresponding fluorescence emission spectra were measured. Infection with the WT cyanophage led to a rapid and substantial decrease in the chlorophyll *a* (Chl) peak with time (Fig. 2a, Extended Data Fig. 2a). This decline was first obvious at 3-h post-infection (Fig. 2a), prior to the onset of cell lysis (Fig. 1b). In contrast, infection by the Δ*nblA* knockout cyanophage led to a more gradual decrease in Chl fluorescence (Fig. 2a, Extended Data Fig. 2a). These findings indicate that energy transfer from PE to Chl was rapidly interrupted during infection with the WT phage, but considerably less so with the Δ*nblA* mutant phage.

**Fig. 2.**
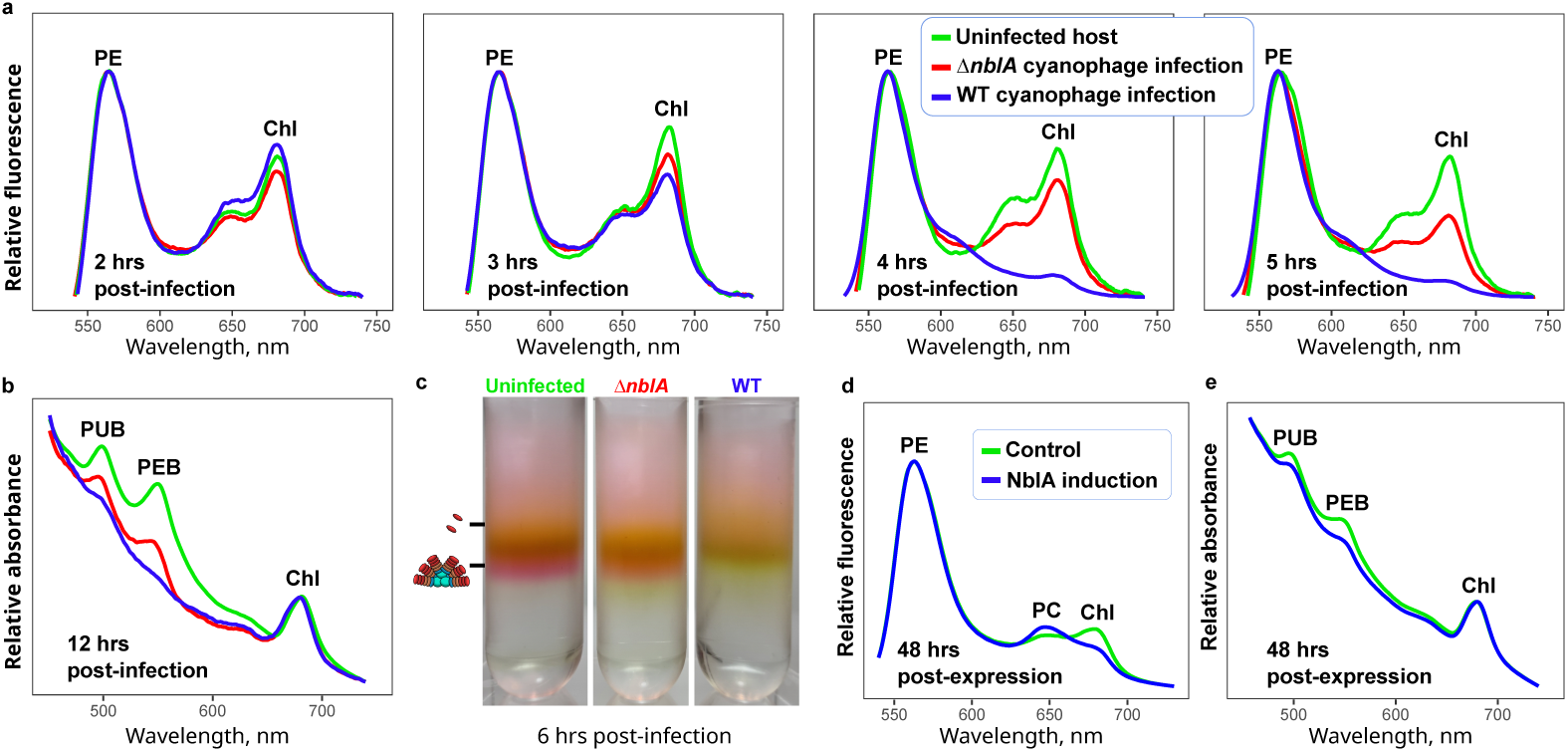
Effect of cyanophage NblA on the cyanobacterial host photosynthetic antennae. **a**, Fluorescence emission spectra, **b**, absorbance spectra, and **c**, isolated phycobilisomes, of uninfected *Synechococcus* sp. strain WH8109 cells (green), and WH8109 infected by the WT (blue) or Δ*nblA* mutant (red) S-TIP37 cyanophages at different time points post-infection. In **c**, the lower density (upper) bands are of disassembled phycobilisome subunits, and the higher density (lower) bands are of intact phycobilisomes (see Extended Data Fig. 3). **d**, Fluorescence, and **e**, absorbance spectra of WH8109 cells 48-h after induced expression of the cyanophage NblA protein (blue) compared to the non-induced control (green). Fluorescence and absorbance spectra are representatives of 5 biological replicates. Averages and standard deviations are shown in Extended Data Fig. 2. Measurements were taken at room temperature, and peaks of Chl, PC and PE peaks are indicated. Fluorescence measurements were normalized on the PE peak and absorbance measurements were normalized on the Chl peak. In **a** and **d**, the PE peak at 560 nm is made up of PEI and PEII, to which the PEB and PUB chromophores bind.

Whole-cell absorbance spectra and proteomic analysis provide a means to assess the levels of host phycobilisome subunits during infection. At 12 hours post-infection with the WT cyanophage, we observed a flattening of the peaks corresponding to both the PUB and PEB chromophores, which are attached to the PEI and PEII proteins. A less prominent decrease in these peaks was observed during infection with the Δ*nblA* mutant (Fig. 2b), suggestive of NblA degradation of the PE proteins by the WT phage. Proteomic analysis during infection with the WT phage revealed significantly lower levels of the α subunits of PEII, PEI and PC rods (CpeA, MpeA and RpcA, respectively), as well as two rod linker proteins (MpeC and MpeE) relative to infection with the Δ*nblA* mutant at 4-h post-infection (Suppl. Data File 1). A 10%-30% reduction was observed for some of the β subunits, but this was not statistically significant. In contrast, three linker proteins closer to the core^35^ that connect PC to APC and both PC and APC to the thylakoid membrane (ApcC, CpcG1, CpcG2) and one rod linker protein (CpeC) had higher protein levels during infection with the WT relative to the Δ*nblA* mutant (Suppl. Data File 1). This may be indicative of attempts by the host to refurbish the phycobilisome core structure in response to the reduction in active phycobilisome rods. Combined, these findings provide further evidence for the role of cyanophage NblA in phycobilisome disassembly and degradation.

Next, we wished to determine the fate of the antenna complex during infection. To achieve this, we isolated phycobilisomes from host cells six hours post-infection and compared them to those in uninfected cells. Fractionation of phycobilisomes on a sucrose gradient followed by fluorescence spectra analysis (Fig. 2c) allowed us to differentiate between assembled (lower, high density, pink bands) and disassembled complexes (upper, low density, orange bands) (see Extended Data Fig. 3). While intact and disassembled phycobilisomes were apparent in uninfected host cells, only disassembled phycobilisomes were present in cells infected by the WT cyanophage (Fig. 2c). Cells infected by the Δ*nblA* mutant cyanophage were composed of both intact and degraded phycobilisomes (Fig. 2c), indicative of only partial phycobilisome disassembly.

The findings above are consistent with a role for NblA in disassembly and degradation of the phycobilisome during infection with the WT cyanophage. However, these findings could also be due, at least partially, to the beginning of cell lysis from 4-h onwards during infection with the WT phage. To more directly assess the impact of phage NblA on host phycobilisomes without the confounding myriad effects of phage infection, we ectopically expressed the phage NblA in host cells without infection. First, we measured whole cell fluorescence spectra which yielded a decline in the energy transfer from PE to Chl, seen by a reduction in the Chl fluorescence peak after excitation at PUB, relative to cells without induction of NblA expression (Fig. 2d). This was, however, to a lesser extent than that found during infection with the WT phage (Fig. 2a). Second, ectopic expression of the phage NblA resulted in some reduction in the PEB and PUB absorbance peaks (Fig. 2e). Third, we assessed whether the cyanophage NblA could restore phycobilisome disassembly during infection with the Δ*nblA* cyanophage mutant. Indeed, NblA expression led to a decrease in the Chl fluorescence peak when excited at PUB, declining to levels similar to those during infection with the WT cyanophage by 3-4-h post-infection (right panels of Extended Data Fig. 4). Note that the partial decline in Chl fluorescence at 4-h post-infection in the control without induction (left panels of Extended Data Fig. 4) is likely caused by leaky expression in this inducible system^33^. Taken together, our findings suggest a clear effect of the cyanophage NblA on energy transfer from PE to Chl, consistent with a role in phycobilisome disassembly and degradation during infection.

Next, we assessed the protein targets of the cyanophage NblA protein by following the production of new protein cleavages in host cells ectopically expressing the viral NblA. For this, we analyzed proteins with newly produced N-termini using mass spectrometry^36^ at 24 and 48 h post-induction (at which time maximum protein induction occurs^33^). Many new cleavages were observed (Suppl. Data File 3). Among these, 209 cleavage events in 107 proteins were markedly more abundant (≥2-fold) following viral NblA expression, including 81 cleavages in 59 proteins at 24 hours and 138 cleavages in 68 proteins at 48 hours post-induction (Fig. 3b, Suppl. Data File 3). These include several cleavages of phycobilisome subunits (Fig. 3a, Extended Data Fig. 5). At 24 h post-induction, four phycobiliproteins were cleaved (Fig. 3b,c); the peripheral PEII disk α subunit (MpeA), the phycocyanin β subunit RpcB, the rod linker MpeE and the ApcE linker responsible for energy transfer from the APC core to the chlorophylls of the photosynthetic reaction centers in the thylakoid membranes^9^. At 48 h post-induction, many of the new proteolytic cleavages were of phycobilisome subunits (Fig. 3a,b). These included the same three PEII, PEI and PC rod disk α subunits that displayed low levels during infection with the WT phage (MpeA, CpeA and RpcA) (see above) as well as the three PE rod disk β subunits (MpeB, CpeB and RpcB), one core protein (ApcB) and three linker proteins connecting between disks in the rods (CpeC, MpeD and MpeE) (Fig. 3b,c, Suppl. Data File 3). These proteolytic cleavages spanned almost all phycobilisome rod proteins, and ApcB of the phycobilisome core (Fig. 3c). These results provide further evidence that the cyanophage NblA protein leads to the degradation and disassembly of the cyanobacterial phycobilisome antenna complex. They also suggest that the cyanophage NblA directs disassembly in a stepwise fashion, initially to disconnect the host phycobilisome from the thylakoid membrane, and to degrade peripheral PEII disks, followed by cleavage of the more internal PEI and PC disks. This is similar to the process known for cyanobacterial NblA during nutrient stress in which phycobilisome degradation begins at the peripheral rod disks and moves inwards towards the phycobilisome core^11^. Intriguingly, Ma et al.^37^ reported that infection by a cyanophage lacking a detectable *nblA* gene leads to the release of PE peptides, suggesting that cyanophages without *nblA* may have an alternative mechanism for degrading outer PE rods.

**Fig. 3.**
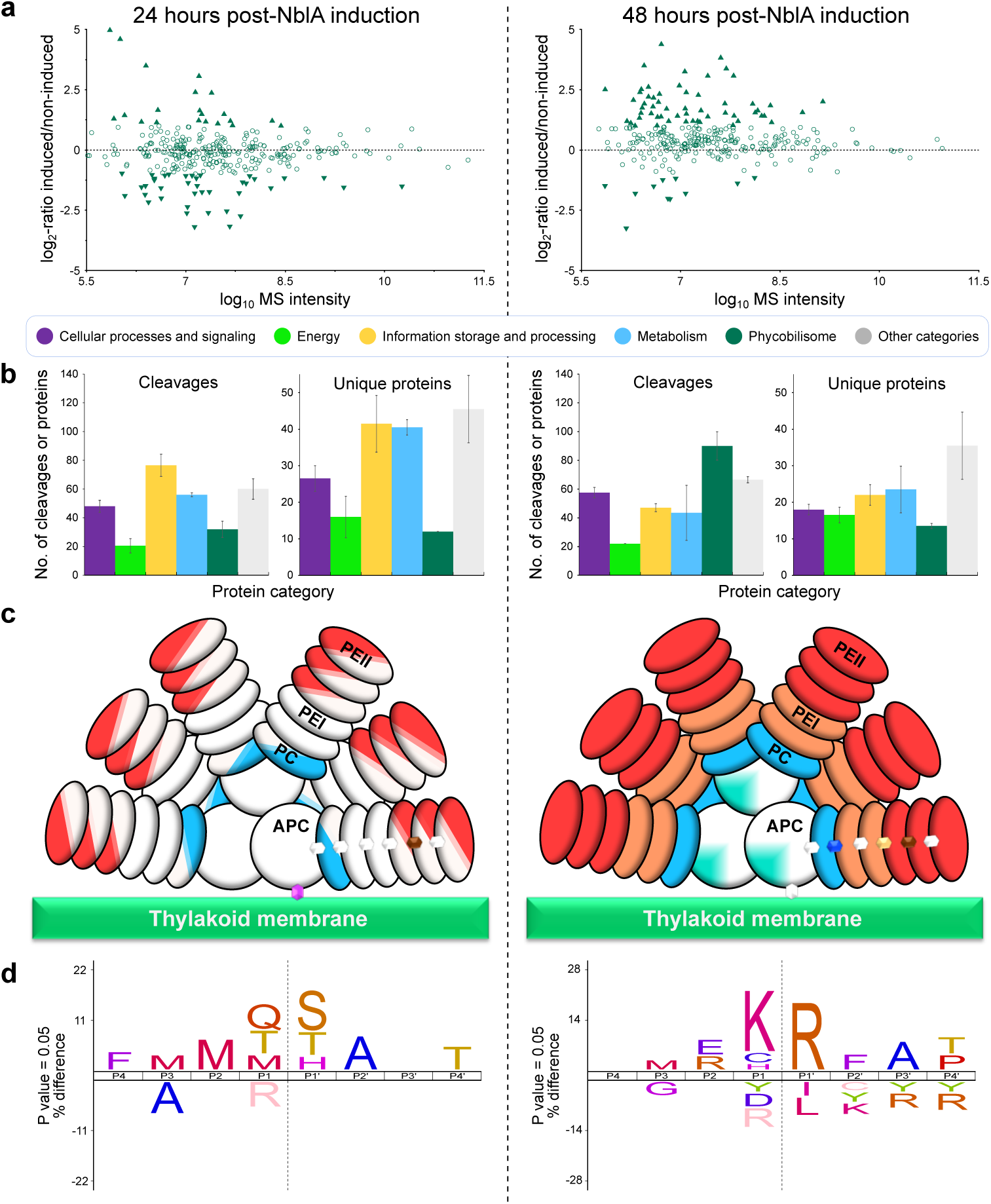
Proteolytic patterns in host cells upon ectopic expression of the cyanophage NblA. Proteomic analysis of newly cleaved proteins after expression of the cyanophage NblA in *Synechococcus* sp. strain WH8109 cells for 24 h (left panels) and 48 h (right panels), as compared to non-induced cells, *n*=2. **a**, MA plots showing the abundance ratios of neo-N-terminal peptides from phycobilisome proteins plotted against their MS signal intensity. This signal intensity serves as an indicator of the peptides’ overall abundances. Filled triangles indicate neo-N-terminal peptides from proteins that show absolute log_2_-ratios ≥1 in both replicates. Triangles with upward-pointing and downward-pointing apexes represent peptides more and less abundant, respectively, upon NblA ectopic expression relative to the control. **b**, The numbers of cleavages and unique cleaved proteins in each category that exhibited a significant abundance change upon NblA expression. Proteins are categorized according to a modified classification of COG functional groups (see Methods), with photosynthesis-related proteins other than phycobilisome proteins included in the “Energy” group. **c**, Phycobilisome cartoons showing proteolytic cleavages in the phycobilisome subunits making up the discs. Colors indicate phycobiliproteins and linker proteins that underwent cleavages in APC (turquoise), PC (blue), PEI (orange) and PEII (red), the linker proteins between the discs, the core and the thylakoid membrane (various colors). Semi-colored discs represent cleavages detected only in the PEII alpha subunits and PC beta subunits. Quarter-colored cores represent cleavages detected only in the APC beta subunits. Fully colored discs indicate cleavages detected in both the alpha and beta subunits. **d**, Preferred cleavage motifs upon ectopic expression of the cyanophage NblA. The amino acid sequence logo is for neo-N-terminal peptides that were identified only after S-TIP37 NblA expression and that had abundance ratios that were at least 2-fold higher in the NblA expressing cells. Cleavage sites are marked with a dashed line.

To our surprise, ectopic expression of the cyanophage NblA led to proteolytic processing of additional photosynthesis-related proteins beyond phycobilisome proteins (Extended Data Fig. 5, Suppl. Data File 3). These include those related to PSI (PsaC, PsaK), the PSII oxygen evolution complex (PsbO, PbsU), and carbon fixation (CcmK2, CsoS2). This suggests that the cyanophage NblA negatively impacts host photosynthesis through proteolytic degradation not only of phycobilisome components but also of reaction center proteins and proteins needed for carbon fixation. Moreover, a set of proteins with functions other than those related to photosynthesis were also cleaved as a result of expression of the cyanophage NblA (Extended Data Fig. 5). These include proteins involved in translation, with the majority of cleavages being in ribosomal proteins at 24 h post-induction (Fig. 3a,b). Translation initiation and elongation factors were also cleaved 48 h post-induction (Fig. 3a,b, Suppl. Data File 3). Other key cellular proteins cleaved include proteins involved in carbon metabolism, a subunit of the cellular RNA polymerase (RpoC2), the cell division protein FtsZ, as well as porin and transporter proteins (Fig. 3b, Suppl. Data File 3). It is not known whether host NblA proteins also direct the degradation of these proteins. Cleavage of these proteins would thus disrupt multiple cellular processes during infection, in addition to photosynthesis.

In freshwater cyanobacteria, NblA proteins function by binding to phycobilisome subunits as well as to the ClpC chaperone responsible for recruiting the protease that induces proteolytic degradation of the phycobilisome^11,38^. Since proteolytic enzymes often have preferred cleavage sites, we investigated cleavage sites after ectopic expression of the cyanophage NblA in the *Synechococcus* host (Fig. 3d). At 24 h post-induction, we detected cleavages primarily between medium-sized, polar or partially polar residues (Met, Gln, Thr) and small, polar or nucleophilic residues (Ser, Thr, His), while at 48 h post-induction an additional cleavage site was detected between Lys and Arg residues (Fig. 3d), providing an explanation for the additional cleaved proteins at 48 h post-induction. This could be a result of NblA recruiting different proteases or that the different substrates are cleaved by the same proteolytic system (presumably the Clp protease complex) that has broad cleavage specificity.

## Cyanophage *nblA* is common in the oceans

T7-like cyanophages are very abundant in the oceans^31,39^. However, the prevalence of the *nblA* gene in this group is largely unknown. Therefore, to understand the environmental significance of our findings, we first assessed the distribution of *nblA* genes in assembled genomes of both isolated and environmental T7-like cyanophages. Our search revealed that 46% of the complete non-redundant T7-like cyanophage genomes code for *nblA* genes (Extended Data Fig. 6). *nblA*-encoding cyanophages have been isolated on picocyanobacterial hosts from the three main lineages: marine *Synechococcus* subcluster 5.1 (*Parasynechococcus*^40^ in the broad sense), *Synechococcus* subcluster 5.2 (*Cyanobium* and related genera) and *Prochlorococcus* (the *Prochlorococcus* collective) (Fig. 4), even though the vast majority of the latter do not have phycobilisomes as their light-harvesting antennae^21,23^. We also found this gene to be present in a rare prophage residing in a *Synechococcus* subcluster 5.1 genome, AG-670-B23^41^. Thus, the cyanophage *nblA* gene is quite common in the genomes of T7-like cyanophages, infecting both *Synechococcus* and *Prochlorococcus*.

**Fig. 4.**
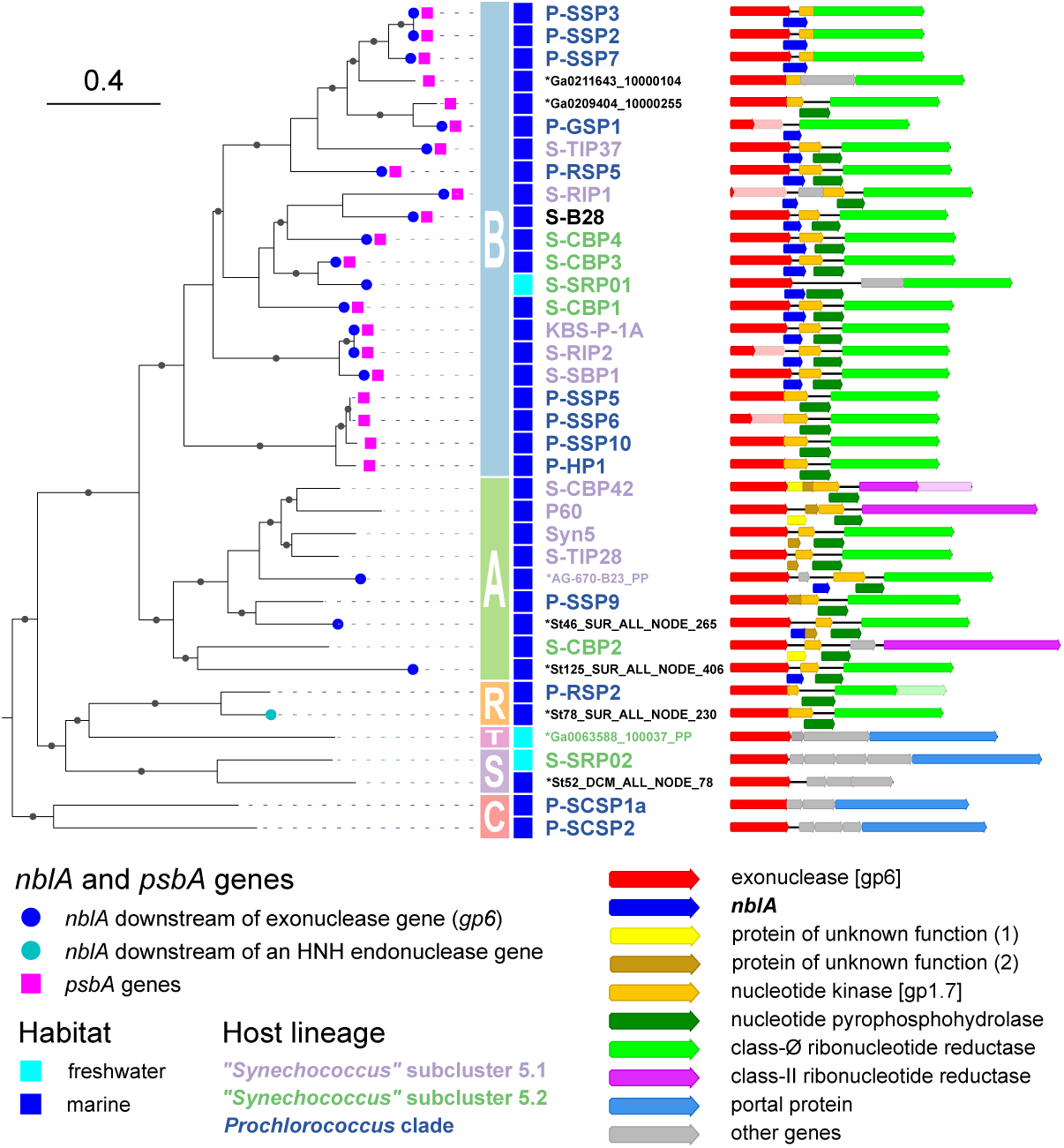
Distribution of *nblA* genes among isolated T7-like cyanophages and selected environmental phage genomes. Phylogenetic tree is based on concatenation of nine core genes (see Methods), is outgroup-rooted and was pruned to include the chosen genomes (see Extended Data Figs. 5 and Suppl. Data Files 4, 5 for the complete phylogeny of the cyanophages and the associated metadata). The genomes are subdivided into previously delineated clades A, B and C, as well as the newly defined minor clades: R (P-RSP2-like phages), S (S-SRP02-like phages) and T (no isolated representatives). For each representative, the genomic region containing the exonuclease gene and downstream of it, is shown. Frequently-appearing orthologous genes are indicated with color. Asterisks indicate environmental genomes. Solid circles mark branches with ultrafast bootstrap support values ≥95.

A close examination showed two types of *nblA* genes with differential distribution among the phylogenetic clades of T7-like cyanophages (Fig. 4 and Extended Data Figs. 6,7). *nblA* genes of one type are dominant and are present in clade B cyanophages (to which S-TIP37 belongs) and to a lesser extent in clade A. These *nblA* genes are consistently located downstream of the exonuclease gene and frequently have up to a 110-nt overlap with it. *nblA* genes of another type are restricted to some P-RSP2-like cyanophages (designated here as clade R) and are located downstream of an HNH endonuclease gene. Neither *nblA* gene type was found in the genomes of cyanophages from the newly discovered clade C (Fig. 4).

To assess the relationships between NblA proteins from the T7-like cyanophages and those of their hosts, we performed a targeted search for NblA homologs among marine picocyanobacteria. To our surprise, we found *nblA* genes not only in marine *Synechococcus*, but also in many *Prochlorococcus* low-light-adapted ecotypes (Extended Data Fig. 7,8). While marine *Synechococcus* typically possesses several *nblA* genes, with e.g. *Synechococcus* sp. strain WH8109 having five intact genes and a pseudogene, *Prochlorococcus* strains have a single *nblA* gene. These findings indicate that *nblA* genes are also found in cyanobacteria that do not possess phycobilisomes as well as in the cyanophages that infect them.

NblAs from the marine picocyanobacteria are substantially divergent from the well-characterized freshwater cyanobacterial NblAs (as exemplified by the chromosomal NblA from *Nostoc* sp. strain PCC 7120) with respect to sequence, yet their predicted structures preserve the classical dimeric NblA architecture (Extended Data Fig. 8). However, NblA from the T7-like cyanophages demonstrate higher overall sequence similarity to the well-characterized cyanobacterial NblAs rather than to those of their hosts (Extended Data Fig. 8). The T7-like cyanophage NblAs cluster together in a phylogenetic network, suggesting that they represent a monophyletic group (see Extended Data Fig. 7). Together these findings suggest that the acquisition of *nblA* by T7-like cyanophages occurred in the distant evolutionary past. This is in contrast to *nblA* from freshwater cyanophages infecting *Microcystis* and *Planktothrix* which have high similarity to *nblA* genes from the corresponding cyanobacterial groups^6,8,27^.

We now turned to assessing the relative abundance of T7-like cyanophages carrying the *nblA* gene in the oceans. We made use of the consistent genomic association of the dominant *nblA* type with the exonuclease gene, a core gene found in all T7-like cyanophages sequenced so far^43,44^, to interrogate the read data from the Global Ocean Viromes (GOV) metagenomic dataset. This allowed us to determine both the presence or absence of the *nblA* gene and the relative abundance of the different clades from the exonuclease gene. Our quantification strategy yielded general distribution patterns of T7-like cyanophages similar to previously reported studies. Cyanophages belonging to clades B and C dominated the global oceans^31,43^, while clade A and R phages were relatively rare. The vast majority of cyanophages from clade B possess the *nblA* gene (Fig. 5), with ∼72% of those at the surface, and 89% of those deeper in the photic zone at the deep chlorophyll maximum (DCM), carrying the gene. The proportion of *nblA*-encoding cyanophages among the less abundant clade A group was much lower, averaging ∼24% at the surface and 10% at the DCM. In accordance with the distribution of the *nblA* genes in complete genomes, none of the clade C and R cyanophages had *nblA* genes downstream of the exonuclease gene. Since the *nblA* gene is located in a different genomic position in clade R, our quantification method could not be applied to them. Overall ∼35% and 65% of the T7-like cyanophages (from clades A, B and C) coded for the *nblA* gene in the surface and at the DCM, respectively. These findings indicate that *nblA* is widespread amongst the abundant clade B cyanophages both at the surface and deeper in the photic zone.

**Fig. 5.**
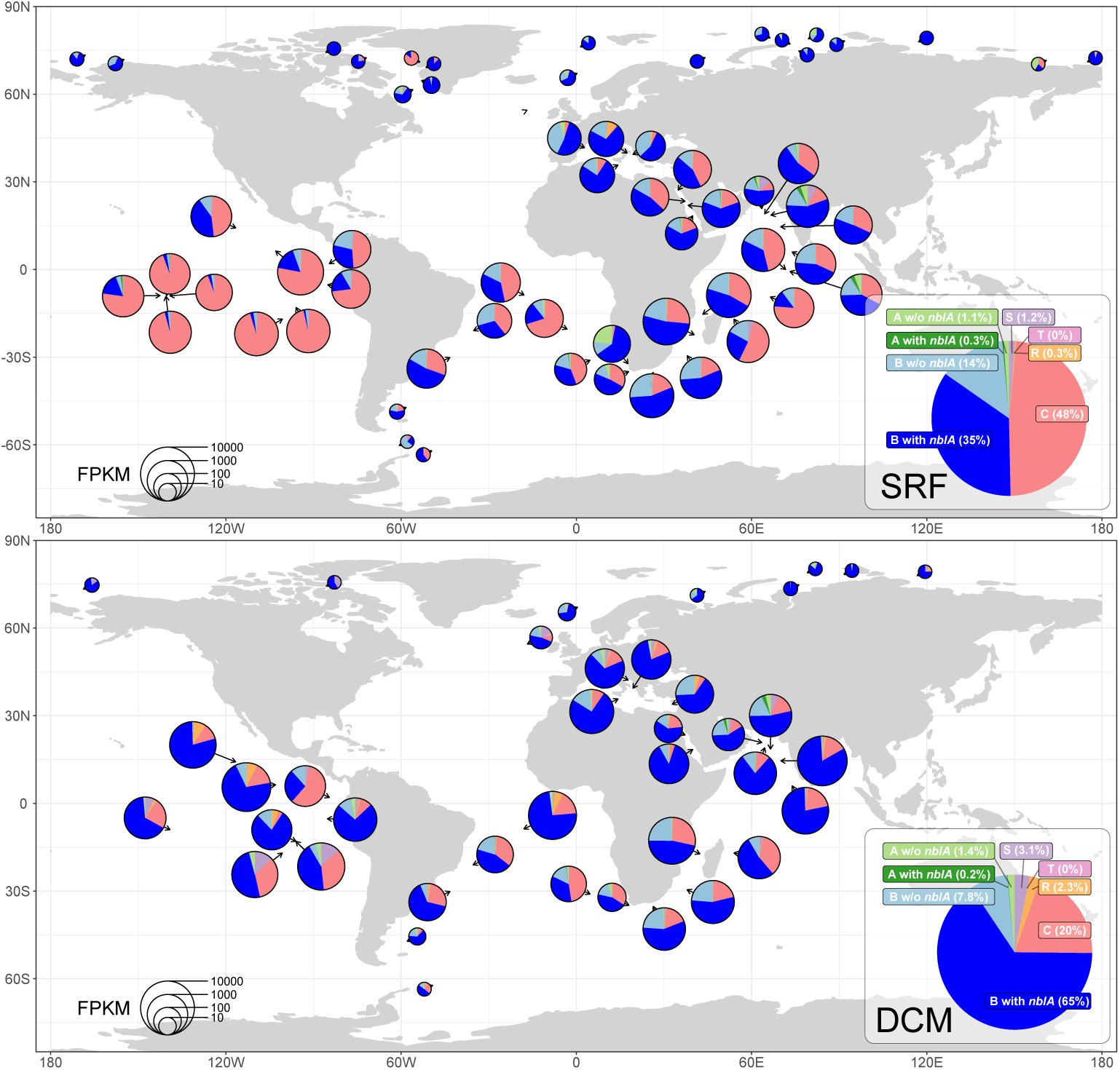
Global distribution of T7-like cyanophages with and without *nblA* genes. **a**, Surface waters. **b**, Deep chlorophyll maximum (DCM). Quantifications are based on fragments per kilobase per million total reads (FPKM) abundances for exonuclease genes with and without downstream *nblA* genes and assigned to one of the five clades of T7-like cyanophages. Raw data and exonuclease-containing contigs for read recruitment were retrieved from the Global Ocean Viromes 2.0 dataset (GOV 2.0)^54^. The insets show the relative abundances of the different clades of cyanophages across the data (based on the FPKM values added together) and serve as the legend for the colors on the maps.

Among the T7-like cyanophage isolates that code for the *nblA* gene, half infect marine *Synechococcus* strains while the rest infect phycobilisome-less *Prochlorococcus* strains (Fig. 4). Both groups of cyanobacteria are prevalent in surface waters, whereas *Prochlorococcus* is considerably more abundant than marine *Synechococcus* at the DCM^21,45^. Thus, we anticipate that many of the clade B phages at the DCM infect *Prochlorococcus*. Since the vast majority of *Prochlorococcus* lineages have a different type of light harvesting antenna to the phycobilisomes of most cyanobacteria^21–23^, we expect that NblA of the phages infecting *Prochlorococcus* direct the degradation of other proteins. This possibility is supported by the finding that multiple low-light-adapted *Prochlorococcus* ecotypes code for an *nblA* gene (Extended Data Fig. 7), even though they do not have phycobilisomes. Furthermore, our proteomics data indicate that NblA-directed degradation is not restricted to phycobilisome proteins, even for a clade B phage that infects *Synechococcus* (Fig. 3a,b, Extended Data Fig. 5). Thus, other photosynthesis-related proteins and proteins involved in other cellular processes, are likely targeted by many *nblA*-encoding cyanophages deep in the photic zone of the oceans.

Cyanobacteria acclimate to low light to maximize their ability to harvest light^41^. This is achieved through increasing the number and size of their antenna complexes^46–48^ as well as the number of their photosystems^49^. As such, more cyanobacterial resources are bound up in the photosynthetic apparatus at depth. Thus, we propose that the ability of cyanophages to degrade the photosynthetic apparatus deep in the photic zone would provide a considerable advantage through the release of amino acids for use in cyanophage progeny production and may explain why relatively more cyanophages at depth carry the *nblA* gene.

Interestingly, the general distribution of the dominant *nblA* type in T7-like cyanophages parallels that for the *psbA* gene (coding for the PSII reaction center protein D1). Both genes are present in the majority of clade B cyanophages, sporadically in clade A, and are absent from clade C. However, they do not always co-occur (see Fig. 4 and Extended Data Fig. 6, see also^28^). The *nblA* gene is more common in T7-like cyanophages than the *psbA* gene, with 36% of the complete genomes (including environmentally assembled genomes) across all clades coding for *psbA* compared to 46% for *nblA* (see above). Since the *psbA* gene is considered to be one of the most common auxiliary metabolic genes in cyanophage genomes^2,3,28,43,44^, our results suggest that the cyanophage-encoded *nblA* gene is very common in oceanic environments as well.

An intriguing aspect of T7-like cyanophages that carry both *psbA* and *nblA* is that *psbA* is thought to enable continued photosynthetic energy production during infection^3,50,51^, while *nblA* degrades the complex that harvests the light funneled to the photosystems (this study). How can these potentially conflicting functions for two different auxiliary metabolic genes in the same cyanophage be reconciled? We propose that these genes are most beneficial to the phage under different conditions. The *psbA* gene has been hypothesized to be more important under conditions of high light^52,53^ where more photodamage to the D1 protein is expected. In contrast, *nblA* is likely to be more important to the phage under low light conditions where more of the cell’s resources are invested in building the photosynthetic apparatus (see above). In addition, under high light conditions, degradation of the phycobilisome by cyanophage NblA could serve to reduce photodamage to D1^7^, such that photosynthesis can continue at some level despite less light being harvested by the phycobilisome. Thus, a balance between the activities of these two proteins is likely needed at high light.

The vast abundance of clade B T7-like cyanophages in the oceans coding for the *nblA* gene raises the possibility that they have a global impact on cyanobacterial photosynthesis. This group of cyanophages is considerably more abundant deep in the photic zone than at the surface^31,55^ where a very large percentage of them code for *nblA* (Fig. 5). While it is difficult to quantify the global impact of a single gene, we have performed some rough calculations to estimate the potential effect of viral *nblA* on picocyanobacterial light harvesting. We base our calculation on the findings that the cyanophage gene caused a 50% reduction in PSII photosynthetic performance (Fig. 1e), that 1-15% of marine cyanobacterial cells are infected by T7-like cyanophages at a given time^39,56,57^, and that 35-65% of marine T7-like cyanophages code for *nblA* in different layers of the photic zone (Fig. 5) (see Methods). We estimate that 0.2-1.1% and 1.6-4.9% of cyanobacterial photosynthesis in the surface and DCM layers, respectively, could be impacted by *nblA*. These estimations suggest that, collectively, T7-like cyanophages carrying the *nblA* gene have a global impact, reducing picocyanobacterial photosynthetic light harvesting by 0.2-5% in the upper oceans.

## Concluding remarks

Auxiliary metabolic genes in viruses^58,59^ are a widespread phenomenon, especially in marine cyanophages^1–4^. Previous studies have investigated the putative function of several such genes using *in-vitro* biochemical approaches^60,61^ or ectopic expression^6,13,61,62^. Here we combined a recently developed cyanophage genetic engineering system^14^, together with inducible ectopic expression^33^ and state-of-the-art N-termini proteomics^36^ to directly examine the role of *nblA*, a gene coding for a small proteolysis adaptor, in cyanophage infection and its impact on the host photosynthetic apparatus. While NblA in cyanobacteria serve as a stress response mechanism under nutrient deprivation and mediate the controlled degradation of the phycobilisome photosynthetic antenna^9–11^, we propose that its cyanophage counterpart is important not for cell survival, but for virion synthesis. Our results further reveal that the cyanophage NblA directs degradation not only of host phycobilisomes, but also of a suite of other proteins, including important core photosynthesis and house-keeping proteins. Thus, this small auxiliary metabolic gene is likely to have a large impact on both the host’s light harvesting efficiency, energy production, and on other essential cellular processes.

The implications of the *nblA* gene for the cyanophage are dramatic with an approximately 3-fold shorter infection cycle, suggestive of considerably improved fitness. These findings, together with genomic and metagenomic results of high abundance and widespread distribution of cyanophages carrying the *nblA* gene in the oceans (this study), and in freshwater ecosystems^7,8,13^, suggests that these cyanophages have a negative global effect of up to 5% on the amount of light harvested by oceanic cyanobacteria.

## Methods

### Cyanobacterial growth

S*ynechococcus* sp. strain WH8109 was grown in artificial seawater (ASW) medium^63^, with modifications as described in Lindell et al.^64^. Cultures were grown at 21 °C and at a light intensity of 20 µmol photons m^−2^ s^−1^, under a 14:10 light-dark cycle. Pour-plates were obtained with ASW medium with low-melting point agarose at a final concentration of 0.28% with additional 1 mM of sodium sulfite. Heterotrophic helper strain, *Alteromonas* sp. EZ55, was added to the pour-plate mixture for isolation of *Synechococcus* colonies^65^.

### Quantitative reverse transcriptase real-time PCR (qRT-PCR)

For RNA extraction, a 1 ml culture sample was harvested by centrifugation at 4 °C, 15,000*g* for 15 minutes and the pellets were flash frozen in liquid nitrogen. Cell pellets were thawed and incubated with lysozyme L6876-5G (SIGMA ALDRICH) at a final concentration of 30 mM, and 200 units of RNase inhibitor (Murine BioLabs) for 60 minutes at 37 °C. An equal volume of lysis buffer was added and cell debris were centrifuged at 4 °C, 16,000*g* for 1 minute. Nucleic acids in the supernatant were precipitated with an equal volume of 95% ethanol and centrifuged as described above. RNA wash buffer (500 µl) from Monarch Total RNA Miniprep Kit (NEB #T2010) was added, and the sample centrifuged again for 30 seconds. The supernatant was discarded and the step was repeated. DNase I reaction buffer (5 µl) and 4 units of DNase I from the TURBO DNA-free™ Kit were added to the 45 µl sample and incubated for 10 minutes at 4 °C. Samples were incubated with 0.5 µl of 0.5 M of EDTA, pH 8.0, for 10 minutes at 75 °C.

Total RNA was reverse transcribed into cDNA using the LunaScript® RT SuperMix Kit (New England Biolabs, E3010). The reaction mixture was prepared in a total volume of 20 µl, 4 µl of LunaScript RT SuperMix containing random hexamers, and 16 µl of nuclease-free water, with the RNA sample added to reach the final volume. As a control, a similar mixture was prepared, but without addition of the reverse transcriptase enzyme. The reaction was incubated at 25 °C for 2 minutes, 55 °C for 10 minutes, and 95 °C for 1 minute.

Quantitative PCR (qPCR) reactions were prepared using LightCycler 480 SYBR Green I Master mix from Roche, combined with 0.2 µM of each primer and the *nblA*/*rnpB* DNA template. The reactions were run on a LightCycler 480 Real-Time PCR System. Cycle threshold fluorescence values for each reaction were determined using LightCycler 480 software. To quantify DNA copy numbers, a standard curve was generated by running 10-fold serial dilutions of the template and correlating cycle threshold values to known DNA concentrations. See Suppl. Data File 6.

The genes whose expression levels were analyzed with qPCR were as follows: host *nblA2* gene (Syncc8109_1607, see explanation below, primers 5’-GCGATCAAGCGGTCAATCAAC-3’ and 5’-CTCTCTGCCGCACGTAGAGG-3’), host *rnpB* gene (Syncc8109_0157, primers 5’-CATCGGCGGTGTGTTTCT-3’ and 5’-CAGGCTTGCTGGGT-3’), S-TIP37 *nblA* gene (primers 5’-TTCCCGAGGCAGACAAGAG-3’ and 5’-TAATGGGATGGTGACTCGGC-3’), S-TIP37 DNA polymerase gene (STIP37_17B, primers 5’-TGAGCTACTACGCAACAGGC-3’ and 5’-AGCGCGATCATTCAGGGAAG-3’). The *Synechococcus* sp. strain WH8109 *nblA* gene chosen for qPCR is the one we reported previously^6^, although a more refined remote homology search with hhsearch^66^ using a custom NblA profile, reveals that the genome carries four additional *nblA*-like genes and an *nblA* pseudogene. In order to clarify which homolog has the highest structural similarity to previously characterized proteins and is thus most likely to have a function similar to that of freshwater cyanobacterial NblAs, we obtained the structures for the corresponding monomers with ColabFold^67^ and searched them against the Protein Data Bank with Foldseek^68^. The best match indeed obtained for NblA2 (Syncc8109_1607), with the highest similarity to NblA protein from *Nostoc* sp. PCC 7120^10^ (PDB: 1OJH.E, probability 0.94, TM-score 0.6621).

### Cyanophage infection experiments

Prior to infection, *Synechococcus* sp. strain WH8109 was grown in liquid medium to mid-log growth to ∼1 × 10^8^ cells ml^-1^. Infection experiments were initiated by adding the S-TIP37 cyanophage strains at a multiplicity of infection (MOI) of 5. Infection dynamics were determined by collecting samples at hourly time intervals during the initial six hours, followed by sampling every two hours thereafter. Samples were filtered over a 0.22 µm syringe filter and the filtrate containing free cyanophages was plated to determine the number of infective cyanophages using the plaque assay^69^ in semi-solid pour plates (see above). Statistical analysis of the infection course was performed with segmented linear regression by selecting near-linear ranges in the infection curves and obtaining linear fits with factors time post-inoculation, cyanophage type (wild-type or mutant) and their interaction and replicate as a random effect using lme4 v. 1.1-31^70^. Significance of the model terms was tested with the Anova function from the car package v. 3.1-1^71^. Distributions of the residuals were checked with QQ-plots.

### Spectral measurements

Absorbance spectra were measured at room temperature using a double beam Shimadzu spectrophotometer equipped with deuterium and halogen lamps. Spectral data were obtained at a constant bandpass with a resolution of 1 nm. Absorbance spectra were normalized to chlorophyll (680 nm). Fluorescence emission spectra were collected using the Jobin-Yvon Horiba Spectrofluorometer. Samples were placed in 1 ml micro quartz cuvettes, excited at 495 nm and fluorescence emission measured from 500 nm to 750 nm and normalized to the fluorescence intensity at the emission maximum of PEI (562-564 nm).

Apparent photochemical quantum yield of PSII (QY_p_), or efficiency of light utilization by PSII, was determined by measuring fluorescence emission, after dark adaptation of 15 minutes, with and without the addition of (3-(3,4-dichlorophenyl)-1,1-dimethylurea (DCMU) to a final concentration of 10 µM. DCMU blocks electron transfer between the primary quinone electron acceptor (Q_A_) and the secondary quinone electron acceptor (Q_B_) on the acceptor side of PSII, causing the reaction centers to remain in a closed state^72^. Samples were excited at 495 nm and fluorescence emission was measured from 500 nm to 750 nm. The apparent PSII quantum yield was calculated as described by Schoffman & Keren^73^, from the onset of infection up to ten hours post-infection, at which point all values fell below zero. QY_P_ was calculated as QY_P_ = (QY_F+DCMU_ -QY_F-DCMU_) / QY_F+DCMU_. Absorbance-corrected fluorescence quantum yield (QY_F_) was calculated as QY_F_= *F*/*f*_495_ where F is the integrated area under the fluorescence emission spectrum, and *f*_495_ is the fraction of incident light actually absorbed at 495 nm (*f*_495_ = 1 - 10^-*A*_495_; *A*_495_ is the measured absorbance at 495 nm).

Quantification of the absorption levels of the different pigments was performed by subtracting the background with the package baseline v. 1.3-4^74^ using the ‘modpolyfit’ method, identifying the peaks of PUB and PEB and chlorophyll and scaling the intensities to the intensity of the chlorophyll peak. Differences between the uninfected cells and cells infected with the wild-type and mutant cyanophages 12 h post-infection were analyzed with linear models and Tukey’s post-hoc tests. Fluorescence emission spectra were scaled to the maximum intensity (intensity of the PE peak) to obtain the relative emission intensity of the chlorophyll peak. Since the fluorescence measurements were performed repeatedly for the same batches, a mixed model was built to analyze the relative fluorescence of chlorophyll as a function of infection (uninfected cells and two types of cyanophages), time post-infection, their interaction taking into account the random effect of batch with lmer4. Distributions of the residuals were checked with QQ-plots. Anova function from the car package was used to obtain analysis of deviance tables and *p*-values for the model terms, Tukey’s test was performed as post-hoc in both cases.

### Phycobilisome extraction

*Synechococcus* sp. strain WH8109 cells were harvested by centrifugation at 6000*g* for 10 minutes at 4 °C, 6 hours post-infection and kept at -80 °C. Pellets were thawed at room temperature, resuspended with 7.5 M K-Phosphate, pH 7.5, and homogenized. Cells were disrupted with a microfluidizer (Microfluidics Corporation HC-2000) at 80 psi. The supernatant was collected after centrifugation at 20,000*g*, for 30 minutes at 4 °C and loaded onto a 0.25-1.25 M linear sucrose gradient in 7.5 M K-Phosphate buffer, pH 7.5. Samples were spun down in an ultracentrifuge at 170,000*g* for 18 hours at 4 °C. Fractions were collected with a syringe and analyzed by fluorometry at 77K and spectrophotometry as described above.

### Construction of S-TIP37 *nblA* deletion mutant

The open reading frame (ORF) encoding the *nblA* gene is 237 bp long (NCBI accession MH540083.1, located at 12,467-12,706 nt) and was previously identified by us in the genome of S-TIP37 based on a match to our custom NblA protein profile^6^. Profile-profile searches with hhsearch via the HHpred server^75^ for this protein yield high-scoring matches to cyanobacterial NblA sequences and Pfam NblA profile (PF04485). Analogously, folding the protein with ColabFold^67^ produces a typical dimeric structure of NblA with the two monomers composed of two helices each and unstructured terminal regions (see Ext. Data Fig. 7).

The *nblA* mutant was produced following Shitrit et al^14^. Since phages gain random mutations readily, we constructed two independent S-TIP37 Δ*nblA* mutant cyanophages to ensure that the observed phenotype was directly related to the lack of the *nblA* gene. Briefly, the *nblA* ORF was used to prepare a construct containing a recombination template for viral homologous recombination. We replaced 137 bp of the *nblA* gene from position 12,572 to 12,707 with a tag of 23 bp. Since the *nblA* gene has an overlap of 94 nt with the upstream exonuclease ORF *exo*, the region chosen for the deletion covered the part of *nblA* not overlapping this gene. However, this region covered the part of *nblA* overlapping a downstream ORF coding for a protein homologous to gp1.7, a nucleotide kinase, of the T7 *Escherichia coli* phage, resulting in deletion of 41 nt at the 5’ end of the gene. (This short gene is under-annotated in T7-like cyanophage genomes and was not originally annotated in S-TIP37). The construct was cloned into the replicative pRL-proCAT^14^ plasmid and conjugated into *Synechococcus* sp. strain WH8109^14,76^. The strain expressing the recombination template was infected with the WT S-TIP37. The lysate was filtered over a 0.22 µm syringe filter to remove cell debris. The presence of recombinant phages was verified by PCR with one primer for the inserted TAG sequence and one in the phage genome; 5’-TGGTGATCAGACCGATGGG-3’ (forward) and 5’-GAGCTCATAGCAAAGAAGACGTC-3’ (reverse).

Enrichment and PCR screening for recombinant cyanophages was carried out in 96-well plates containing the *Synechococcus* host^14^. Wells containing recombinant cyanophages were filtered and plated on semi-solid medium. Single phages were plaque purified twice and the whole genome of each purified mutant phage clone was sequenced alongside the corresponding WT. Illumina MiSeq trimmed read pairs were received and the genomes were assembled *de novo* with Spades v. 3.14.1^77^. Both Δ*nblA* mutants had additional mutations in their genomes compared to their corresponding wild-type genomes (see Suppl. Data File 7 for the genome sequences). Non-synonymous mutations in the first mutant relative to the first WT had a T>C substitution at position 11,142 (position according to reference genome MH540083.1) leading to a V208A mutation in the gene for DNA polymerase nucleotidyl transferase subunit (accession AXF42115.1) and an A>C substitution at position 37,714 in gene 42 coding for a putative 2OG-Fe(II) oxygenase (accession AXF42102.1) leading to a K34T change. The second Δ*nblA* mutant had two non-synonymous differences compared to its wild type in gene 40 coding for a putative tail fiber protein (accession AXF42100.1): an A>C substitution at position 36,856 in the wild-type genome leading to a D454A change (relative to the reference genome) and an A>G substitution at 36,888 in the Δ*nblA* genome leading to a N465D change. Since both Δ*nblA* mutants had additional mutations compared to the wild-type, infection and spectral experiments were performed with both mutants and gave the same results. Results with the second mutant are shown. Rescue and proteomics experiments were performed with the second mutant. The mutant does not express the putative nucleotide kinase. The relevance of this gene to the phenotype was addressed in the rescue experiments through ectopic expression of *nblA* but not the putative kinase (see below section).

### Ectopic expression of the cyanophage *nblA*

A theophylline translational induction system was used for expression of the cyanophage NblA protein in *Synechococcus* sp. strain WH8109^32,33^. A riboswitch sequence that theophylline binds to and the full length S-TIP37 *nblA* gene were cloned into the pRL-proCAT replicative plasmid downstream of the *rnpB* promoter following Tahan et al^33^. and was conjugated into *Synechococcus* sp. strain WH8109^14,76^. The construct consists of the regions from 12,467 to 12,703 nt relative to the reference S-TIP37 genome (GenBank accession MH540083.1) and six additional amino acids that were added at the NblA C-terminus (Gly, Ser, Tyr, Ser, Val, Thr) due to the cloning process. Since the *nblA* gene overlaps two neighboring genes, the insert contains the last 95 nt of the 3’ region of the putative exonuclease gene and the first 37 nt of the 5’ region of the putative nucleotide kinase gene. The latter corresponds to 12 out of 94 amino acids of the protein and proteomic analysis verified that this protein was not present in the induction experiments. As such, rescue experiments reinstated expression of NblA but not the putative nucleotide kinase.

Expression experiments were carried out with the exponentially growing WH8109 conjugant. Theophylline (0.3 mM), dissolved in ASW, was added to induce translation of the NblA protein. Expression of proteins under this theophylline-inducible system is leaky, with differences between induced and non-induced expression detected from 7.5 h after addition of theophylline that increased until 48 h post-induction^33^. For rescue experiments theophylline was added 8 h prior to infection with cyanophages. For assessing the effect of NblA on *Synechococcus* sp. strain WH8109 protein cleavage, samples were collected 24 and 48 h post-induction. Induced samples were compared to non-induced samples. Each expression experiment was accompanied by a control construct, carrying the pRL-proCAT plasmid containing the riboswitch but lacking a downstream gene.

### Proteomic analyses

#### Protein extraction

Cyanobacteria were harvested and washed three times using 50 mM of HEPES, pH 7.5. After the final centrifugation the cyanobacterial pellet was resuspended with 8 M guanidine hydrochloride (GuHCl), 100 mM HEPES, pH 7.5, and was heated at 95 °C for 10 min. Following this, the samples were sonicated using VialTweeter (Hielscher) at maximum amplitude, 70% cycle time for 5 min, then heated for another 5 min at 95 °C to ensure maximal extraction and denaturation. Cellular debris were pelleted at 18,000*g* for 10 min, and the clear supernatant containing proteins was transferred to a new tube. Protein concentrations of each sample were measured with BCA, prior to splitting each sample taking 10 µg for total proteome analysis and 50 µg for N-terminome analysis.

#### Total proteomics sample preparation

Proteins were reduced with 5 mM DL-dithiothreitol (DTT) at 65 °C for 30 min then cooled to room temperature (RT) and alkylated with 12.5 mM chloroacetamide (CAA) for 30 min in the dark. Guanidinium concentration was diluted to 1 M using 100 mM 4-(2-hydroxyethyl)-1-piperazineethanesulfonic acid (HEPES), pH 8, and trypsin (Promega, V115A) was added at 1:100 (w/w) ratio for overnight at 37 °C. Next, trypsin was quenched with 1% formic acid and the samples were desalted using C18 stage tips (Empore 66883-U) before subjecting them to mass spectrometry (MS) analysis.

#### N-termini proteomics sample preparation

N-terminal enrichment was done using an optimized Hydrophobic tagging-assisted N-termini enrichment (HYTANE) procedure as described^36^. Proteins were reduced and alkylated with DTT and CAA as described above. Then, different isotopes (heavy and light) of formaldehyde were used to label primary amines in every two comparative samples. The labeling was done with 40 mM formaldehyde and 20 mM of sodium cyanoborohydride at 37 °C overnight. Formaldehyde leftovers were quenched using 100 mM glycine for 1 h at 37 °C before mixing the heavy and light labeled samples together. The samples were diluted to reduce GuHCl concentration to 1 M using 100 mM of HEPES pH 8 before adding trypsin at 1:100 (w/w) and overnight incubation at 37 °C. The trypsin was then quenched with 1% (final) of formic acid, and the samples were subjected to desalting using OASIS-HLB columns (Waters). The elution was done using 60% of acetonitrile, 0.1% formic acid and the samples were dried using a speed-vac. The peptides were resuspended using 100 µl of 100 mM HEPES pH 7 and had undecanal solution (20 mg/ml prepared in ethanol) added to them at w/w ratio of 1:50 (protein:undecanal), followed by the addition of sodium cyanoborohydride at a final concentration of 20 mM. Undecanal tagging was done for 2 hours at 50 °C while renewing the sodium cyanoborohydride after 1 h. The samples were then centrifuged at 16,000*g* for 5 min and the supernatant was transferred to a new tube before drying in a speed vac. The dried samples were resuspended in 500 µl of 2% acetonitrile, 0.1% formic acid and subjected again to OASIS-HLB column. Elution was done using 60% of acetonitrile, 0.1% formic acid and the samples were dried in a speed-vac before resuspending again and subjecting them to MS.

#### LC-MS analysis

Desalted samples were subjected to LC-MS analysis using an Orbitrap Exploris 480 coupled with an EvoSep One HPLC. Samples were introduced onto the EvoTip, which was then washed twice with 20 μl of 0.1% formic acid. The washed peptides remained wet, maintained by applying 150 μl of 0.1% formic acid atop of the EvoTip until MS analysis. The samples loaded on the EvoTips underwent chromatographic separation on a 15 cm × 150 μm analytical column, filled with 1.9 μm C18 beads (EV1106). Peptides were separated over an 88-minute gradient according to the manufacturer’s standard method. Full MS scans acquired in positive ion mode, scanning from 300 to 1800 m/z, were recorded at a resolution of 120,000, by a data-dependent mode selecting the top 20 ions with high energy collisional dissociation (HCD) fragmentation ion at a resolution of 17,500 and with dynamic exclusion enabled.

#### MS data analysis

For total proteomics, data analysis was conducted using FragPipe v. 22.0 (https://fragpipe.nesvilab.org/) using DDA+ mode^78^ with the default Label-Free Quantification Match Between Runs (LFQ-MBR) settings. The search enzyme was set to trypsin, allowing up to two missed cleavages. Variable modifications included methionine oxidation and protein N-terminal acetylation, while carbamidomethylation of cysteine was set as a fixed modification. For N-terminomics data, raw files were first converted to MzML format and then analyzed using the Trans-Proteomic Pipeline version 6.3^79^ with the Comet search engine v. 2023_01 rev2. The search enzyme was set to Semi-ArgC, allowing up to two missed cleavages. Variable modifications included methionine oxidation and the mass difference between heavy and light dimethylation at lysine residues or the N-termini of peptides. Fixed modifications included carbamidomethylation of cysteine and light dimethylation at lysine residues or peptides’ N-termini.

Relative peptide quantifications were performed using XPRESS, with parameters set to mass tolerance of 20 ppm, a minimum of three chromatogram points for quantitation, and the number of isotopic peaks to sum set to zero. Post-analysis table creation, cleavage motif extraction, and ratio normalization were performed using an in-house script^80^. The search database included all *Synechococcus* sp. strain WH8109 proteins in Uniprot (Taxon ID: 166314, both reviewed and unreviewed) in addition to the vector added genes and standard contaminant proteins. For viral infection samples, this database was expanded to also include all *Synechococcus* T7-like phage S-TIP37 (Taxon ID:1332145) protein sequences from Uniprot and the NblA sequence. In the analysis of the N-terminomics data, we considered three classes of blocked N-terminal peptides: ORF N-termini, neo-N-termini generated by internal proteolysis (SemiN1), and proteolysis-adjacent peptides truncated before the first arginine (SemiN2). Only NblA-induced cleavage events that were identified in both biological replicates and showed >2-fold abundance increases in NblA-expressing cells compared to the control in each replicate were considered for downstream analysis. To bin host proteins into functional groups, we utilized the COG functional category assignments for *Synechococcus* sp. strain WH8109 proteins in the eggNOG database v. 5.0^81^. The proteins were assigned to one of the broad functions based on a modified classification of COG functional groups as follows: phycobilisome components — Phycobilisome; other photosynthesis-related genes and other proteins assigned to COG functional category C (energy production and conversion) — Energy; COG functional categories J, A, K, L and B — Information Storage and Processing; COG functional categories D, Y, V, T, M, N, Z, W, U and O — Cellular Processes and Signaling; COG functional categories G, E, F, H, I, P and Q — Metabolism; COG functional categories R, S and proteins assigned to multiple categories — Other.

Phage proteins were classified into functional categories based on the functions of the majority of genes in three genomic and expression clusters^29,44^. Statistical analysis of the dynamics of abundance of the different functional groups of phage proteins in the total proteomics data as a function of time post-infection onset was performed as follows. For each gene cluster cumulative MS LFQ intensities were used as a response variable in a linear model with time post-infection and infection type (by WT or ΔnblA mutant) and their interaction as predictor terms. Distributions of the residuals were checked with QQ-plots. Detailed results of the statistical analysis are provided in Suppl. Data File 2.

### Bioinformatic analyses

#### Analysis of T7-like cyanophage genomes

Genomes of cultured T7-like cyanophages and outgroup phages (coliphage T7 and pelagiphages HTVC011P and HTVC019P) were collected from GenBank. Complete genomes and genomic fragments of environmental T7-like cyanophages and cyanoprophages were obtained from JGI IMG/VR v. 4.1^82^ and Global Ocean Virome v. 2.0 assemblies^54^. To collect genomic fragments containing exonuclease genes with sufficiently long downstream regions, a conserved region in the C-terminal part of the exonuclease (corresponding to residues 86-221 in the exonuclease protein from S-TIP37 [RefSeq accession YP_009807515.1]) was chosen and homologous regions from a reference set of cyanophages were used as queries for tblastn from NCBI blast v. 2.15.0^83^. Fragments were retained when they had matches at least 400 bp long, identity of ≥40%, bit score of ≥400 and at least 500 bp of downstream sequence available. To collect genomes for phylogenetic analysis, out of the contigs with the exonuclease gene, we extracted sequences ≥30 Kbp long and dereplicated them with dRep v. 3.4.5^84^ at the ANImf identity level of 0.95. Genes were predicted with prodigal v. 2.6.3^85^ and phylogeny was reconstructed with phylophlan v. 3.0.2^86^ (diamond v. 2.1.8^87^, MAFFT v. 7.475^88^, trimAl v. 1.4.1^89^, IQ-TREE v. 2.1.2^90^) based on concatenated alignments of nine core genes: primase-helicase, exonuclease, portal protein (head-to-tail adaptor), head assembly protein, major capsid protein, tail tubular proteins A and B, small and large terminase subunits. The genes were extracted by searching with diamond blastp using representative protein sequences as queries from two distantly related cyanophages, P60 and S-SRP02: P60_gp14 (primase-helicase is missing from S-SRP02), P60_gp18 and SSRP02_p034, P60_gp26 and SSRP02_p038, P60_gp27 and SSRP02_p039, P60_gp28 and SSRP02_p040, P60_gp29 and SSRP02_p041, P60_gp30 and SSRP02_p042, P60_gp40 and SSRP02_p011, P60_gp49 and SSRP02_p012, respectively. IQ-TREE was run with “-m LG --alrt 1000 -pers 0.2 -nstop 500” with 1,000 ultrafast bootstrap replicates^91^ and the resulting tree was outgroup-rooted with the ingroup defined as the branch encompassing the known cyanophages. The cyanophage genomes were classified based on the resulting phylogeny into six clades: the previously recognized clades A, B and C, the newly defined clades R (for P-RSP2-like phages), S (S-SRP02-like phages) and T (represented by GOV contig Station52_DCM_ALL_NODE_78). See Suppl. Data File 4 for metadata of the genomes chosen to represent each of the clades. Two outlier genomes were found to cause spurious clustering and were excluded from the phylogenetic analysis: IMGVR_UViG 3300032116_000204 and IMGVR_UViG 3300029337_000223. The incidence of *nblA* and *psbA* genes in the genomes was assessed by searching ORFs (between stop codons) with hmmsearch from HMMER v. 3.4^92^ using a custom NblA profile (see below) and the PsbA profile TIGR01151.1 from NCBI Protein Family Models. The incidence and type of DNA polymerase in the genomes was assessed by searching the ORFs with blastp using the sequences of the exonuclease (YP_009807513.1) and polymerase (YP_009807514.1) subunits of the split polymerase of S-TIP37 with an E-value threshold of 1e-10. A version of the tree focusing on representative genomes as shown in the main text, was obtained by trimming the full tree. The trees were visualized with ggtree v. 3.2.0^93^. The full phylogenetic tree without the outgroups is available in Suppl. Data File 5.

#### Picocyanobacterial genomes

Metadata and genome sequences for picocyanobacteria were downloaded from Cyanorak v. 2^94^ and additional *Prochlorococcus* genomes ^20,25^ were downloaded from Integrated Microbial Genomes & Microbiomes^95^. Genes were predicted with prodigal and GeneMarkS-2 v. 1.14_1.25^96^ with the predictions merged using gffcompare v. 0.12.6^97^.

#### Analysis of NblAs

The NblA HMM profile used to search for *nblA* genes in T7-like cyanophage genomes was obtained from an alignment of the previously released NblA sequences^6^ and manually curated NblA sequences from representative T7-like cyanophages. The searches for the NblA hits were performed without heuristic filters and the results were filtered using an empirically determined full score threshold of 20. The same NblA profile was also initially used to search for *nblA* genes in the protein sequences from picocyanobacteria. Upon discovering that many of their NblAs yield sub-significant hits, we built a separate NblA profile specifically targeting picocyanobacterial NblAs based on the sequences collected in the first round of the search. Notice that due to the high divergence of the NblAs from picocyanobacteria, many of them avoided detection in our previous screen of cyanobacterial genomes using a general NblA protein profile^6^. The two NblA protein profiles are available in Suppl. Data File 10.

A NeighborNet network was constructed to visualize similarity relationships between NblA proteins of different origins. NblAs from T7-like cyanophages and picocyanobacteria were combined with cyanobacterial sequences assigned in UniProt r. 2025_2^98^ to Pfam profile PF04485, as well as selected NblA sequences from non-T7-like cyanophages. To reduce redundancy, the sequences were clustered at 90% identity level with CD-HIT v. 4.8.1^99^. The cluster representatives were aligned with MAFFT in automated mode, and trimAl was used to trim the alignment (strict mode, minimum column block size of 15). The resulting trimmed alignment was used as input to SplitsTree v. 4.17.0^100^ which generated a NeighborNet network based on uncorrected distances. The network was visualized with tanggle v. 1.8.0^101^.

Sequence logos were generated for different groups of NblAs using DiffLogo v. 2.26.0^102^. The original MAFFT alignment was trimmed in a similar way as above but only regions outside of the core NblA domain were trimmed. The fasta file was converted to A3M format (matching columns with <50% gaps) and secondary structure predictions were added with tools from HH-suite v. 3.3.0^66^. For the sequence logos, the remaining gaps were coded as a separate state X.

Representative structures of NblA dimers were predicted with ColabFold v. 1.5.5 (based on AlphaFold2)^67^.

#### Analysis of global distribution of the cyanophage nblAs

For the analysis of the distribution of *nblA* downstream of the exonuclease gene, we took all of the initially collected environmental contigs with T7-cyanophage type exonucleases and extracted regions containing the 3’ end of the exonuclease gene with up to 100 bp upstream and up to 1,000 bp downstream. These fragments were searched against exonuclease sequences from the genomes used for phylogeny using blastx and clades were assigned to fragments with best hits to cyanophages at identity level of ≥60%. The fragments coming from the GOV2.0 assemblies were used for recruitment of GOV raw data with bwa v. 0.7.17^103^. Quantification was performed with featureCounts from subread v. 2.0.6^104^. As the exonuclease and *nblA* genes frequently overlap, we selected strictly defined windows for read quantification instead of ORF boundaries: the exonuclease was represented by the above-mentioned conservative region at the 3’ end of the gene. For the *nblA* gene we chose the location corresponding to the core NblA region based on the hmmsearch matches. Genes assigned to the same cyanophage clade were grouped into metafeatures for quantification and the resulting per-station mapped read pair counts were converted to Fragments Per Kilobase per Million reads (FPKM) values by dividing them by the median feature length (in Kbp) and the total number of the reads per sample (in millions). Sequences used for recruitment and quantification results in tabular format are available in Suppl. Data File 9.

The bioinformatic workflow was implemented in snakemake^105^, python and R.

### Calculations of the ocean-wide impact of viral *nblA* genes on light harvesting by picocyanobacteria

In order to estimate the ocean-wide impact of viral *nblA* genes on light harvesting by picocyanobacteria, we used the following data: A 100% reduction in photosynthetic performance for 50% of the time attributable to the cyanophage *nblA* gene (Fig. 1e); 1-15% of marine cyanobacterial cells are infected by T7-like cyanophages at a given time^39,56,57^; and 35% (surface waters) to 65% (DCM zone) of marine T7-like cyanophages code for the *nblA* gene (Fig. 5). The impact of viral NblA proteins on marine cyanobacterial photosynthesis in surface water was calculated as follows: 50% reduction in photosynthetic light harvesting performance, with 1-6% of cells infected and 35% of T7-like cyanophages carrying *nblA* genes giving an estimated impact of 0.175-1.05% on the photosynthetic performance of the picocyanobacteria. The impact on marine cyanobacterial photosynthesis at the DCM was estimated as follows: 50% reduction in photosynthetic light harvesting performance, with 5-15% of cells infected and 65% of T7-like cyanophages carrying *nblA* genes giving an estimated impact of 1.625-4.875% on the photosynthetic performance of picocyanobacteria.

## Acknowledgments

We thank F. Salama for biophysical guidance, M. Suissa-Szlejf for help in setting up the phycobilisome isolation gradients, R. Edrei for help with spectroscopic measurements, G. Sabehi for discussions on infection experiments, S. Goldin for help with qPCR calculations, S. Larom for technical support, N. Keren and C. Kranzler for discussions on photosynthesis experiments, N. Keren for help in performing spectroscopy, and N. Frankenberg-Dinkel for comments on the manuscript. This work was funded by the European Research Council (ERC Advanced Grant 321647 to O.B. and ERC Consolidator Grant 646868 to D.L.), the Nancy and Stephen Grand Technion Energy Program (GTEP), Israel Science Foundation grants to O.B. (143/18) and O.K. (1515/23), and the Simons Foundation Life Science Award 735081 to D.L. O.B. holds the Louis and Lyra Richmond Chair in Life Sciences and D.L. holds the Dresner Chair in Life Sciences and Medicine.

## Author contributions

O.N. conceived the project, performed viral work, phycobilisome isolation and experimentation, and together with O.B., O.K. and D.L. designed the experiments. I.P. performed phage growth curve experiments. A.R. performed bioinformatic analyses. R.H. and O.K. performed proteomic analyses. D.S., R.T. and D.L. developed the methods for constructing the mutant phage and the cyanobacterial NblA expression strain. O.B. coordinated the project. O.N., A.R., O.B., O.K., and D.L. wrote the paper, which was critically revised and approved by all authors.

## Data availability

All data are available in the main text or the Supplementary Data. Intermediate results of the bioinformatic analyses are available from the GitHub repository https://github.com/BejaLab/autographiviridae. The mass spectrometry proteomics data have been deposited to the ProteomeXchange Consortium via the PRIDE partner repository with the following dataset identifiers: PXD057452 (whole proteome data) and PXD057454 (N-terminomics data) [the reviewers can receive access using the following tokens: ZqvdIvO680Np (project PXD057452) and NpEFI3ZdaRSA (project PXD057454)].

## Code availability

The code used for the bioinformatic analyses is available from the GitHub repository https://github.com/BejaLab/autographiviridae.

## Extended data figures

**Extended Data Fig. 1.**
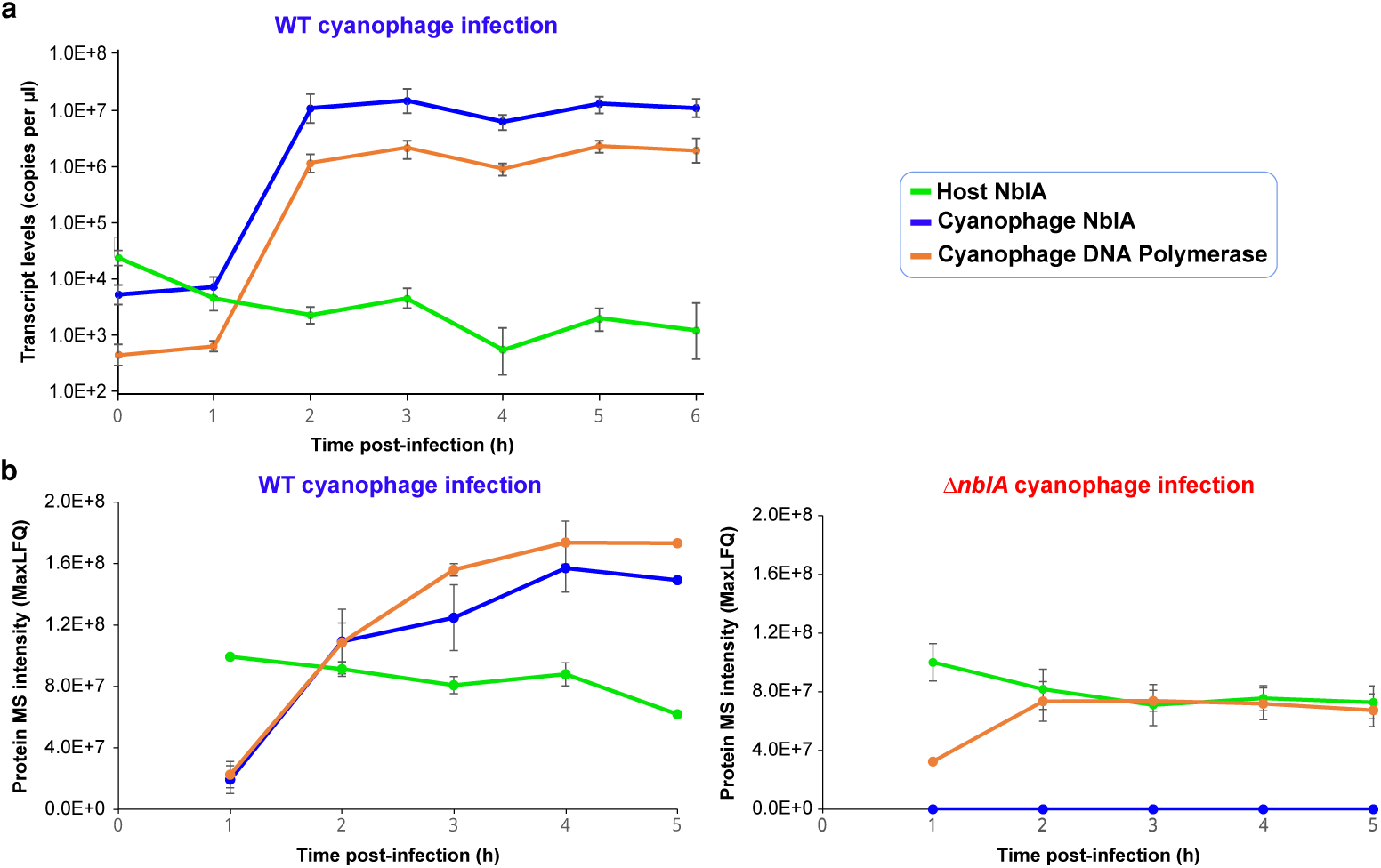
Expression of the cyanophage *nblA* gene during S-TIP37 infection of *Synechococcus* sp. strain WH8109. **a**, Transcript levels of S-TIP37 cyanophage *nblA* and DNA polymerase genes and one of the host *nblA* genes during infection with WT cyanophage determined by qRT-PCR. Transcript levels are normalized to the *rnpB* gene. Among the host *nblA* genes, *nblA2* was chosen as the gene coding for the NblA protein with the highest structural similarity to previously characterized NblAs (see Methods). **b**, Protein mass spectrometry intensity levels of the S-TIP37 cyanophage NblA, DNA polymerase, and host NblA6 proteins during infection with WT (left) or Δ*nblA* mutant (right) cyanophages, determined by LC-MS/MS analysis. NblA6 was the only host NblA protein detected. Data are presented as mean ± standard deviation of three independent biological replicates, *n*=3.

**Extended Data Fig. 2.**
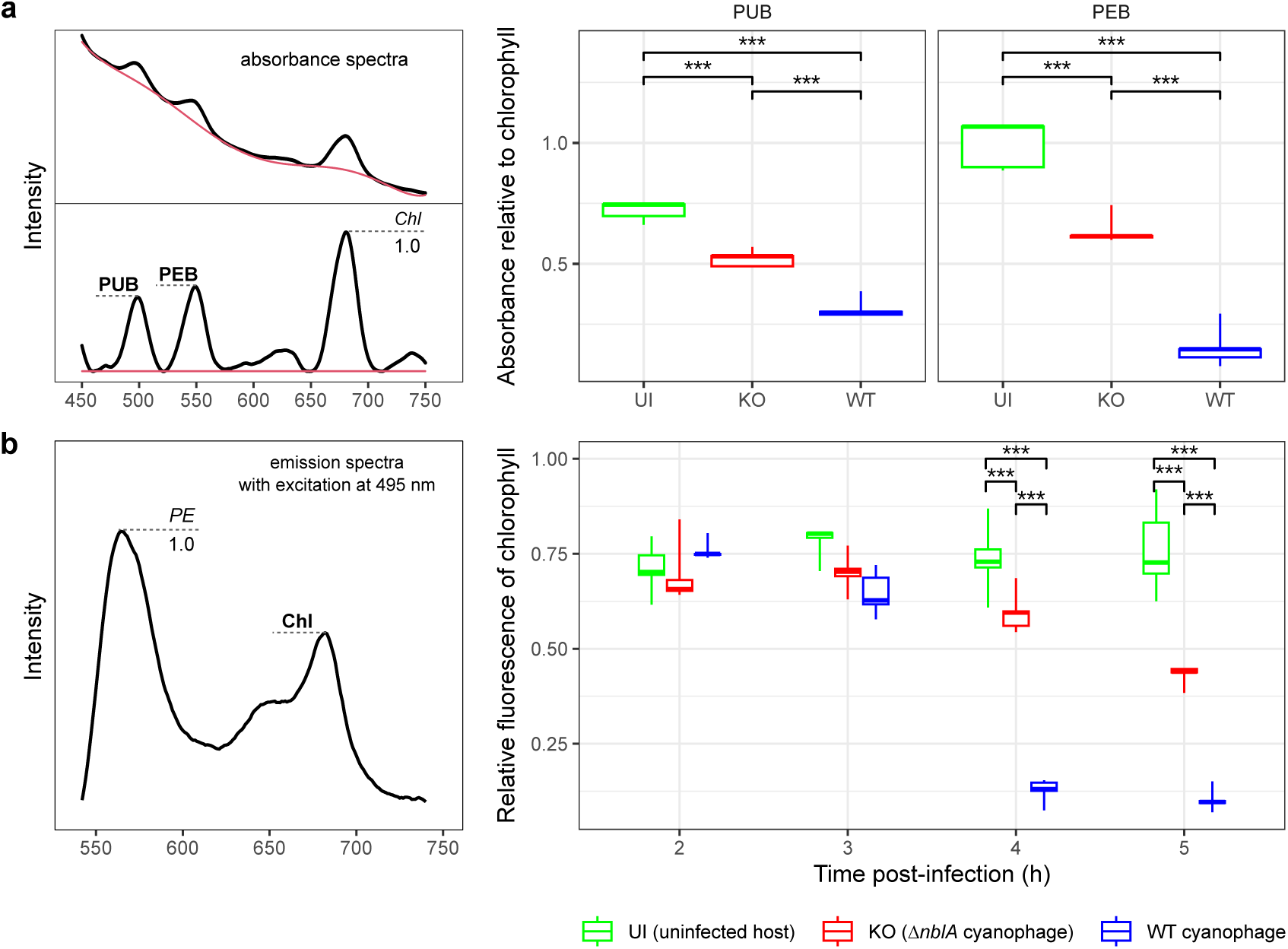
Spectroscopic characteristics of cells infected with wild-type and Δ*nblA* mutant S-TIP37 cyanophages. **a**, Relative absorbance of PUB and PEB at 12 h post-infection. Relative absorbance was determined as follows (see the panel to the left): the absorbance spectra were background-subtracted (red line) and the intensities were scaled to the height of the Chl peak. UI, uninfected, KO, infection with the Δ*nblA* mutant, and WT, infection with the wild-type phage. **b**, Relative chlorophyll fluorescence at different time points during infection. The samples were excited at 495 nm (excitation maximum of PUB), emission spectra were obtained and the relative height of the chlorophyll peak was calculated (see the panel to the left). The PE peak at 560 nm is made up of PEI and PEII, to which the PEB and PUB chromophores bind. In both panels, asterisks denote significant pairs of differences in the Tukey post-hoc test: *** — *p*-value < 0.001 (after FDR adjustment in panel b). Boxplots reflect: minimum-maximum range (vertical line), interquartile range (rectangle) and median (horizontal line). The results are based on five biological replicates of each condition: uninfected cells and cells infected with two cyanophages, Δ*nblA* (KO) and wild-type (WT) phages, sampled repeatedly for measurement. For further results of the statistical analyses refer to Suppl. Data File 2. Representative spectra are shown in Fig. 2a,b.

**Extended Data Fig. 3.**
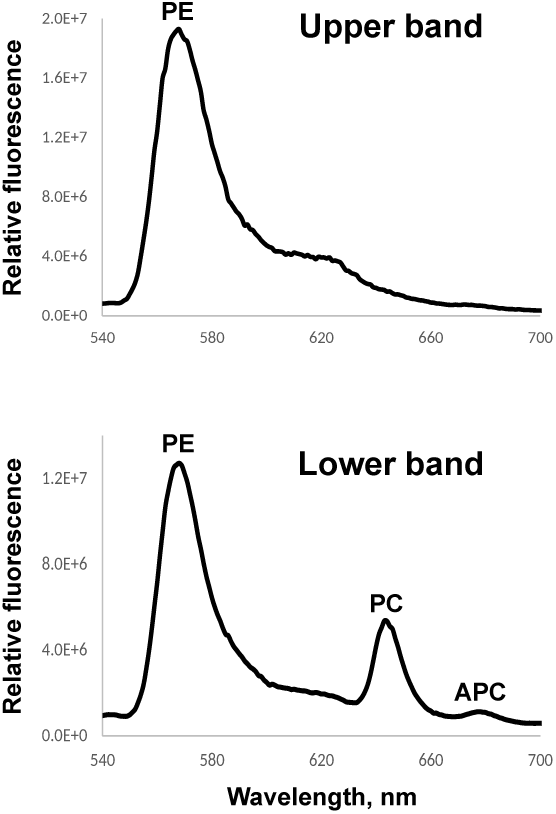
Fluorescence spectral analysis of fractionated phycobilisomes. Fluorescence emission spectra at 77K of fractionated phycobilisomes from uninfected *Synechococcus* sp. strain WH8109. The lower density, upper band (see Fig. 2c) contains disassembled phycobilisome complexes, whereas the higher density, lower band (see Fig. 2c) contains assembled phycobilisome complexes with intact energy transfer from PE to PC and APC.

**Extended Data Fig. 4.**
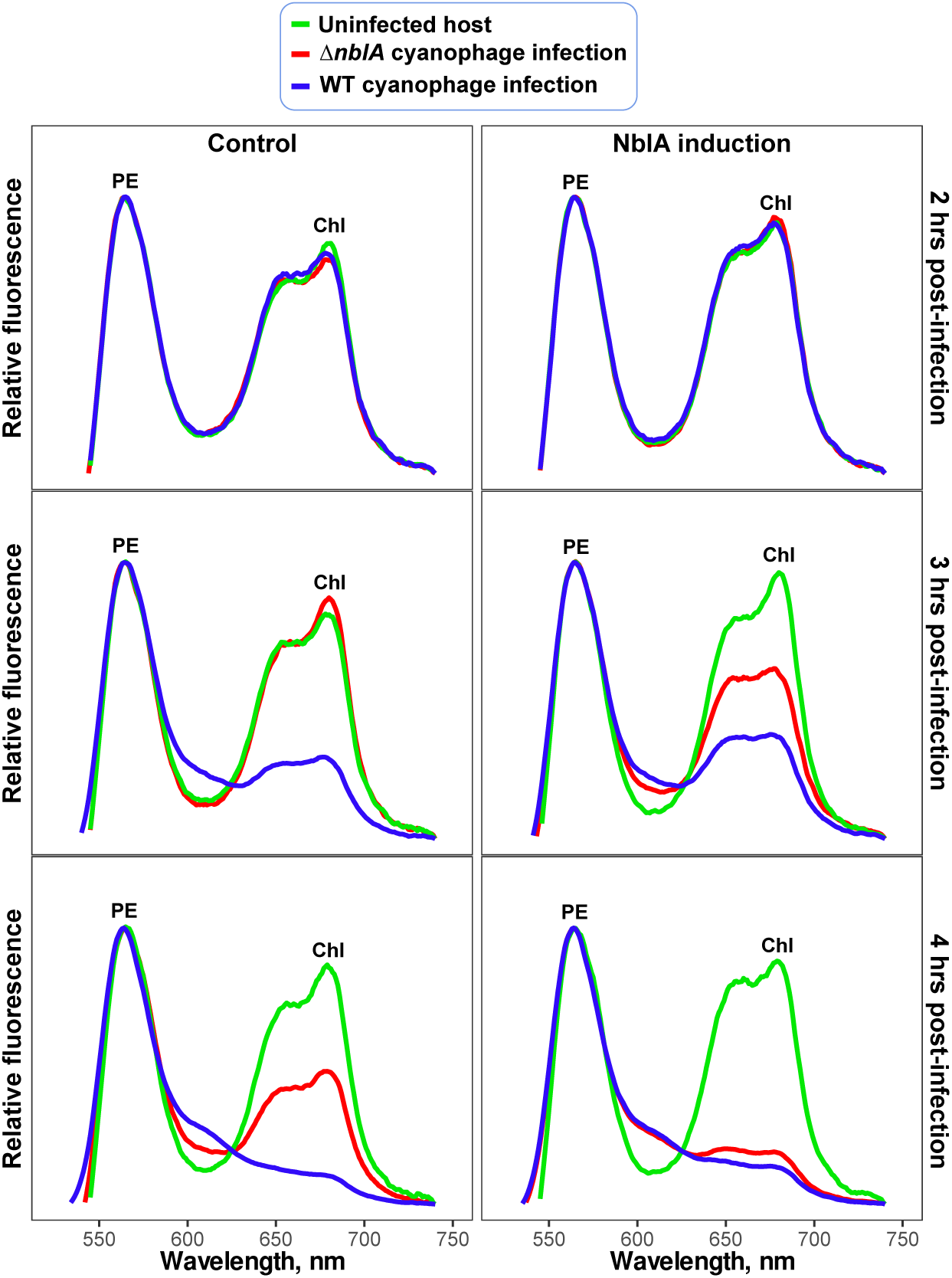
Ectopic expression of cyanophage NblA in *Synechococcus* sp. strain WH8109 during infection. Fluorescence emission spectra of uninfected cells (green), cyanobacteria infected by the WT cyanophage (blue) and by the Δ*nblA* mutant cyanophage (red) at different time points post-infection. Control experiments (the panels to the left) were performed without the addition of the inducer. Ectopic expression was induced in the host 8 h prior to addition of the phages (NblA induction, the panels to the right). Measurements were taken at room temperature and normalized using the PE peak.

**Extended Data Fig. 5.**
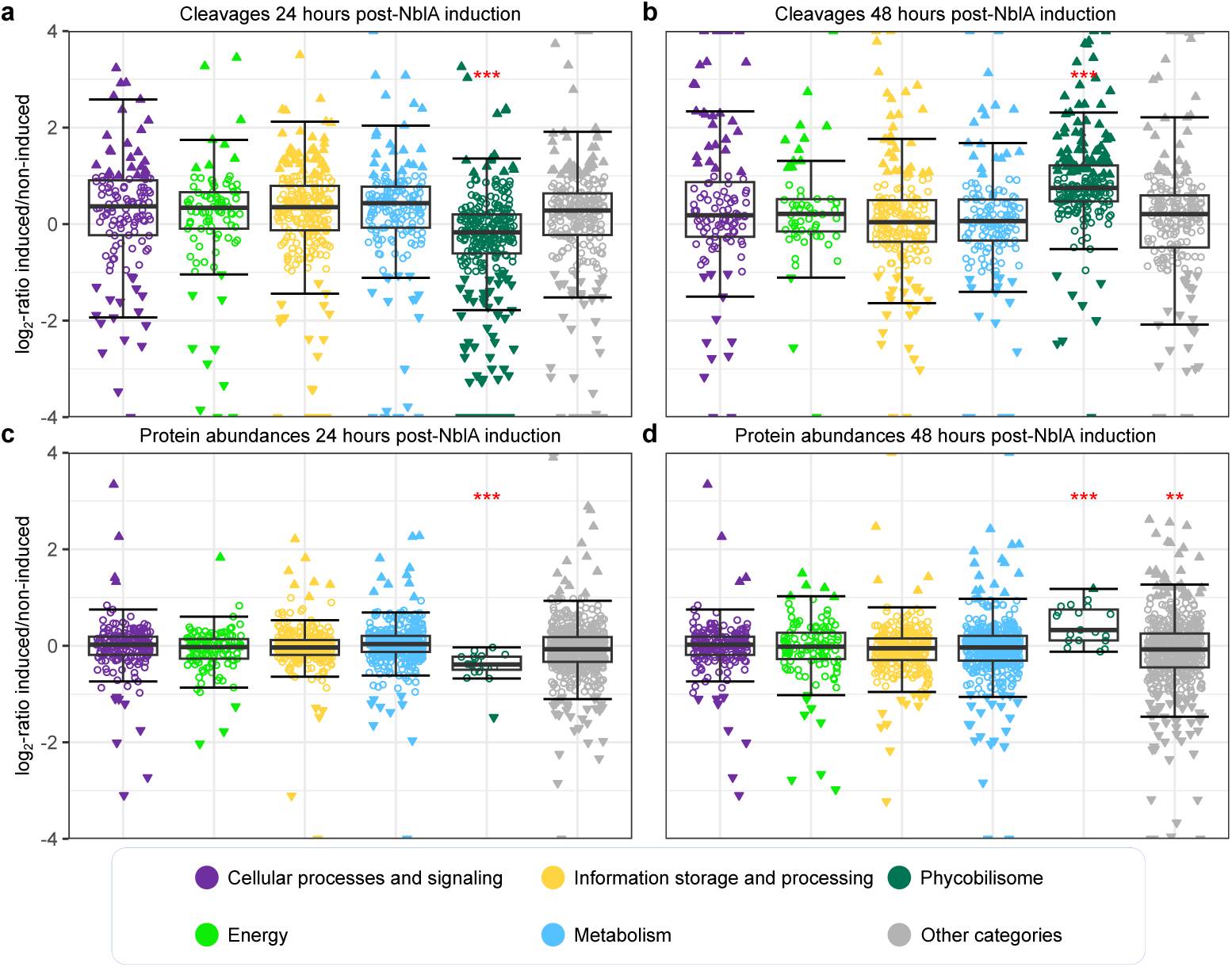
Cleavage and proteome changes upon ectopic expression of cyanophage NblA in *Synechococcus* sp. strain WH8109. Proteomic analysis of cleavage events (**a** and **b**) and total protein abundance changes (**c** and **d**) following ectopic expression of the cyanophage NblA in *Synechococcus* sp. strain WH8109 cells for 24 hours (**a** and **c**) and 48 hours (**b** and **d**), compared to non-induced controls (n = 2). The top panels show the abundance ratios of neo-N-terminal peptides identified by N-terminomics, calculated as the ratio of signal intensities between induced and non-induced samples, reflecting changes in peptide abundance.The bottom panels show the abundance ratios for proteins in the total proteomic analysis based on the intensities of tryptic peptides. Proteins are categorized according to a modified classification of COG functional groups (see Methods). Filled triangles indicate neo-N-terminal peptides and proteins with absolute log_2_-ratios ≥1 in both replicates. Triangles with upward-pointing and downward-pointing apexes represent peptides more and less abundant, respectively, upon NblA ectopic expression relative to the control. Red asterisks indicate significant deviations of the category-wise mean ratios from zero (two-sided one-sample t-test) after FDR adjustment across all comparisons: *** — adjusted p-value ≤ 0.001, ** — adjusted p-value ≤ 0.01.

**Extended Data Fig. 6.**
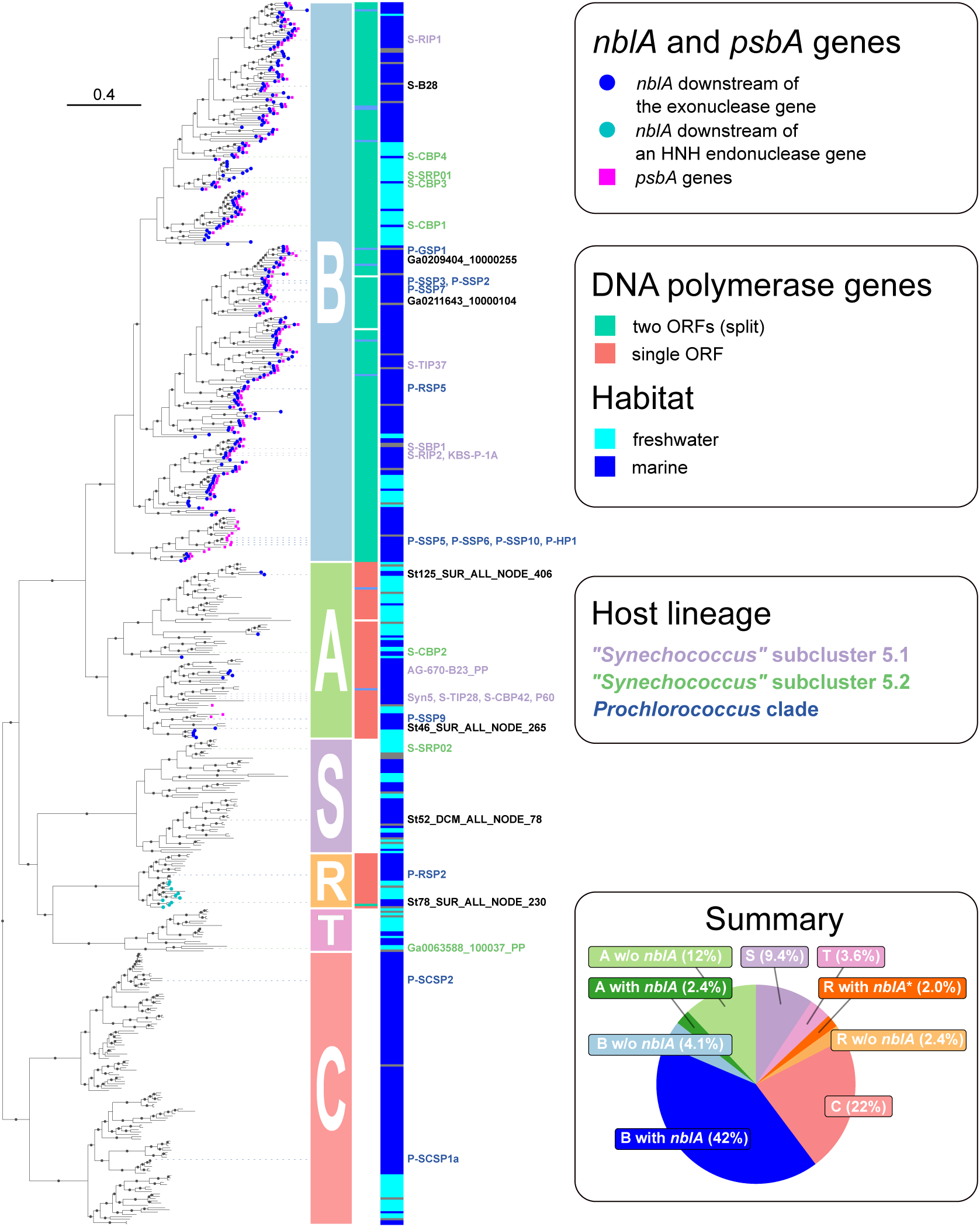
Phylogeny and distribution of *nblA* genes among T7-like cyanophages. Phylogenetic tree was built based on concatenation of nine core genes coding for: primase-helicase, exonuclease, portal protein (head-tail adaptor), head assembly, major capsid protein, tail tubular proteins A and B, and small and large terminase subunits. Phage genomes used in the phylogeny were recruited among cultured T7-like cyanophages and from environmental sources: Global Ocean Virome v. 2 assemblies and JGI IMG/VR v. 4.1. The environmental genomes were recruited by searching for the exonuclease gene, selecting genomes at least 30 kb long, dereplicating them at 0.95 ANImf identity level and requiring the presence of at least eight core genes. The tree was outgroup-rooted by inclusion of coliphage T7, pelagiphage HTVC011P, pelagiphage HTVC019P and environmental phages falling outside of the cyanophage cluster. Phages were categorized into previously recognized clades A (P60-like), B (S-TIP37-like) and C (P-SCSP1a/2-like) and newly-delineated clades R (P-RSP2-like), S (S-SRP02-like) and T (Ga0063588_100037_PP-like, no cultured representatives). Presence and type of DNA polymerase genes is indicated with the clades dominated by phages falling in one of the following categories: no DNA polymerase gene (clades C, S, T), DNA polymerase coded in a single ORF (clades A and R) and a split DNA polymerase gene^106^ with the exonuclease domain coded in a separate ORF (clade B). Clade representatives are colored with labels according to the host’s phylogenetic affiliation if known. Note that each of the three picocyanobacterial lineages represents an assemblage of several genus-level taxa. The incidence of *nblA* (circles) and *psbA* (squares) genes is indicated with symbols at the tips of the corresponding branches and the *nblA* genes of two categories are differentiated: those located downstream of the exonuclease gene (as in S-TIP37) and the *nblA* gene downstream of an HNH endonuclease gene (in a subclade of clade R). No clade C, S or T genomes were found to code for *nblA* and are thus not separated into two groups in the summary pie chart. Solid circles mark branches with ultrafast bootstrap support of ≥95. See Suppl. Data File 7 for the source data.

**Extended Data Fig. 7.**
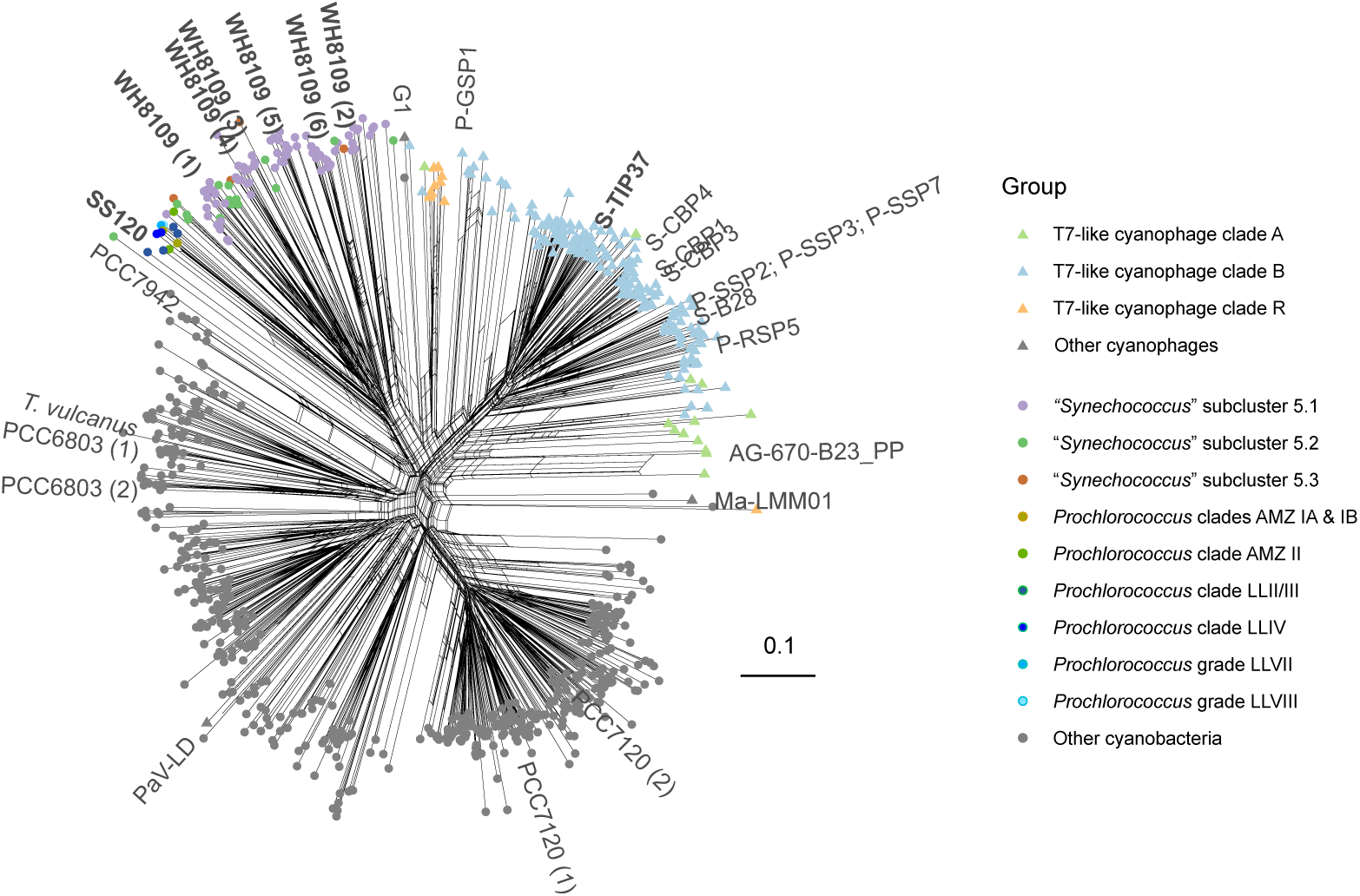
Phylogenetic network of NblA proteins from the T7-like cyanophages, marine picocyanobacteria and other cyanophages and cyanobacteria. The NblA protein sequences were aligned, trimmed and a NeighborNet network was obtained based on uncorrected distances. The proteins are categorized by the corresponding cyanophage or cyanobacterial group. NblA was found in the low-light adapted *Prochlorococcus* ecotypes LLII/LLIII, LLIV, LLVII, LVIII, AMZIA, AMZIB, and AMZII but not LLI and AMZIII nor any of the high-light-adapted ecotypes. NblAs from selected cyanophages and model cyanobacteria with the NblAs from S-TIP37, *Synechococcus* sp. strain WH8109 and *Prochlorococcus* sp. strain SS120 are highlighted in bold. The network and trimmed sequences of cluster representatives in NEXUS format are available in Suppl. Data File 8 and metadata and untrimmed sequences and aligned sequences of the core NblA region are available in Suppl. Data File 11.

**Extended Data Fig. 8.**
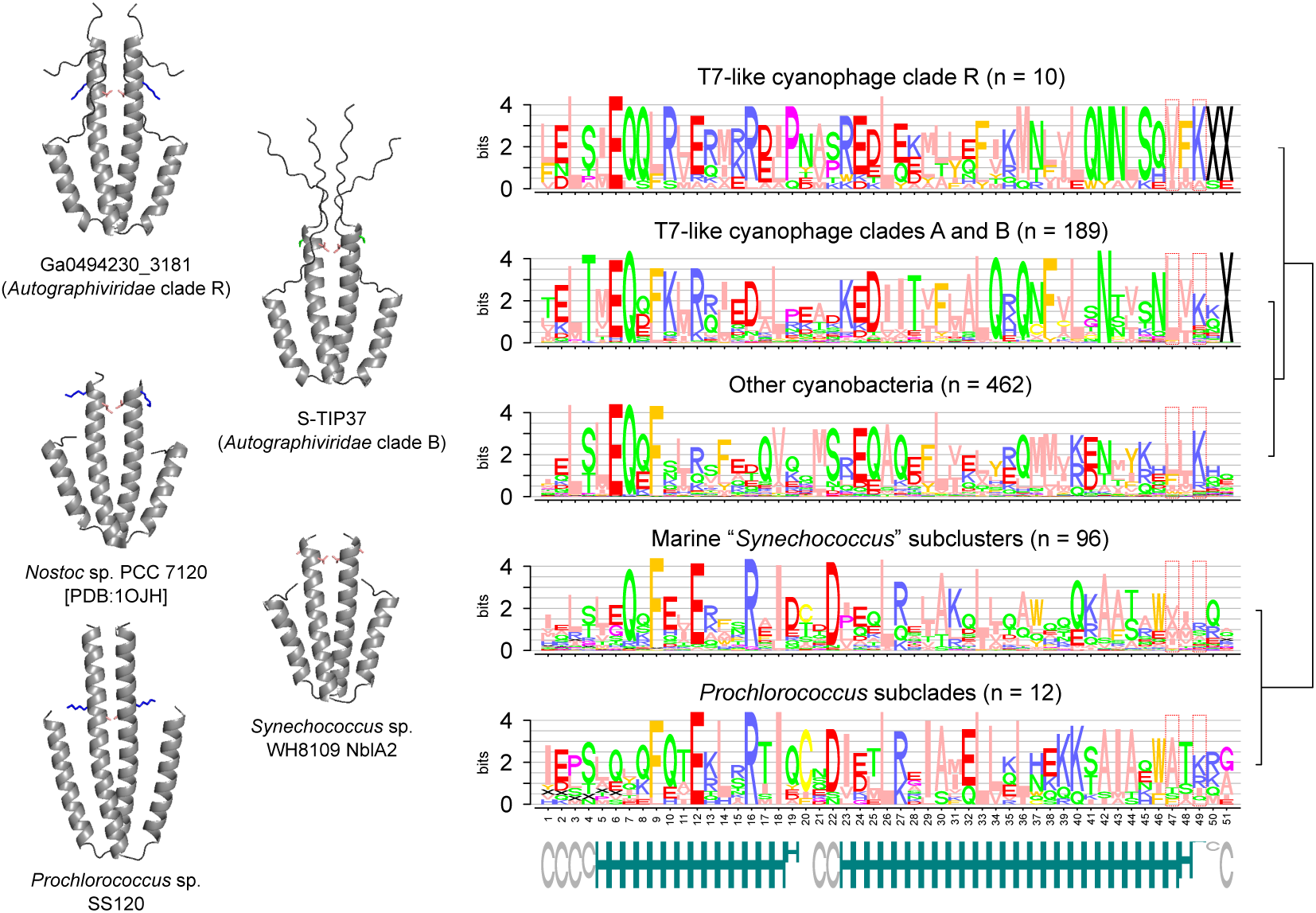
Structures and protein profiles of NblA proteins from T7-like cyanophages and cyanobacteria. Right: dimeric structures of selected representatives of the NblA proteins. All of the structures, except that of the NblA from *Nostoc* sp. strain PCC 7120 [PDB:1OJH^10^] were obtained with ColabFold (AlphaFold2)^67^. Left: sequence logos generated based on a trimmed alignment of clustered NblA sequences. Numbers of cluster representatives per group are provided in parentheses. Secondary structure predictions based on the whole alignment are shown at the bottom (H: helix, C: coil) with the height proportional to the prediction score. Positions known to mediate phycobiliprotein binding^38^ are indicated: protein profile positions 48 (residues L51 in *Nostoc* sp. PCC 7120 and L64 in cyanophage S-TIP37) and 50 (residues K53 in PCC 7120 and T66 in S-TIP37). Note that a variation is found between the NblAs from marine cyanobacteria and their cyanophages at these two positions, which could be indicative of a functional difference between them.

## Supplementary data files

**Supplementary Data File 1.** Mass spectrometry analysis of unlabeled proteins during the infection by S-TIP37. Excel spreadsheet.

**Supplementary Data File 2.** Details of statistical analyses. Excel spreadsheet.

**Supplementary Data File 3.** Whole proteome and Neo N-termini enrichment characterization by mass spectrometry of *Synechococcus* WH8109 due to ectopic expression of the S-TIP37 NblA. Excel spreadsheet.

**Supplementary Data File 4.** Non-redundant T7-like cyanophage genomes used for phylogeny and additional metadata for representative T7-like cyanophage genomes. Excel spreadsheet.

**Supplementary Data File 5.** Rooted and annotated phylogenetic tree of T7-like cyanophages. jtree format.

**Supplementary Data File 6.** qPCR and mass spectrometry data for Extended Data Fig. 1a and b. Excel spreadsheet.

**Supplementary Data File 7.** Genome assemblies of the wild-type and mutant S-TIP37 cyanophages. Fasta format.

**Supplementary Data File 8.** Trimmed amino acid sequences and annotated phylogenetic network of NblA proteins from cyanobacteria, T7-like cyanophages and other selected cyanophages. Nexus format.

**Supplementary Data File 9.** Exonuclease and *nblA* quantification data: sequences used for read recruitment and regions defined for quantification of exonuclease genes and downstream *nblA*, if present. Excel spreadsheet.

**Supplementary Data File 10.** NblA protein profiles: a general profile used for the searches in cyanophage genomes and a picocyanobacterial-specific profile. HMM file.

**Supplementary Data File 11.** NblA proteins collected from T7-like cyanophages and picocyanobacteria and additional NblA sequences used for reference. Excel spreadsheet.

